# DNA-induced conformational changes in SPRTN relieve its auto-inhibitory effect on protease activity

**DOI:** 10.64898/2026.04.29.721615

**Authors:** Wei Song, Joseph A. Newman, Yichen Zhao, Rod Chalk, Christina Redfield, Paul R. Elliot, Kristijan Ramadan

## Abstract

The DNA-dependent metalloprotease SPRTN has emerged as a key enzyme in the proteolysis of DNA-protein crosslinks (DPCs), thereby protecting us against genome instability, accelerated ageing, and cancer. DNA and ubiquitin chains serve as the primary activator and catalyst of SPRTN proteolysis, respectively, but how they promote SPRTN activation and activity remains incompletely understood. To address this question, we developed a highly sensitive multi-turnover fluorescence resonance energy transfer (FRET) assay to monitor SPRTN proteolysis in real time. We found that the auto-cleaved N-terminal SPRTN fragment, comprising the metalloprotease domain (MPD), zinc-binding domain (ZBD), and basic region (BR), is highly stable, enzymatically active, and retains ubiquitin-dependent activation. Interestingly, the MPD alone exhibits basal intrinsic activity that is independent of both DNA activation and ubiquitin avidity effect. We show that ZBD and MPD together exert steric regulation: ZBD maintains MPD in an autoinhibited state, while MPD largely prevents ZBD from binding DNA. BR, together with DNA, is essential to relieve ZBD-mediated inhibition of MPD. Using a site-trapping approach, we demonstrate that the ZBD-BR-DNA trinity induces an open conformation of the SPRTN N-terminus *in cis*, thereby releasing autoinhibition. MPD and BR together restrict the DNA-binding stoichiometry of ZBD, enabling SPRTN to function efficiently in proximity to DNA despite its low abundance *in vivo*. Collectively, our work overturns the long-standing dogma that SPRTN autocleavage inactivates the enzyme and reveals how DNA-induced conformational changes in SPRTN fine-tune its protease activity, providing a prerequisite for subsequent ubiquitin activation and rapid proteolysis of DPCs.

## Introduction

Under a diversity of endogenous inducers, such as reactive oxygen species (ROS), metabolic aldehydes, or external factors such as environmental formaldehyde, chemotherapeutic drugs, UV light, ionising radiation, DNA, and protein could be accidentally and covalently crosslinked to each other in the cells. These undesired DNA-protein crosslinks (DPCs) represent cytotoxic DNA lesions that block replication, repair and transcription, leading to genome instability and cancer^1–6^. The DNA-dependent metalloprotease SPRTN (also known as C1orf124 or DVC1) was identified as the first and essential protease specialised in resolving DPCs^7,8^. Patients deficient in SPRTN develop chromosomal instability, premature ageing and early onset hepatocellular carcinoma^9,10^. Similar findings were recapitulated in SPRTN-haploinsufficient mice and in mice carrying patient variants^11,12^. SPRTN, acting as “molecular scissors”, is essential to remove bulky DPCs at replication forks, preventing fork collapse^7,8^. For example, during replication-coupled interstrand crosslink (ICL) repair, HMCES (5-hydroxymethylcytosine-binding, embryonic stem cell-specific protein) prevents DNA damage by crosslinking to single-stranded DNA (ssDNA) abasic (AP) sites after the CMG complex passes over them. SPRTN is subsequently required to resolve the HMCES-DNA-protein crosslink (HMCES-DPC)^13^. In addition to its role in replication-coupled DPC repair, SPRTN also participates in global-genome DPC repair, which is regulated by SUMO-targeted ubiquitination^14^. A recent study additionally addressed that DPC accumulation in mitotsis caused by SPRTN deficiency results in the leaking of damaged DNA into the cytoplasm. The damaged DNA is subsequently sensed by the cGAS-STING pathway for inflammatory sigalling transdution^11^.

The regulation of SPRTN protease activity has been relatively well studied from multiple angles *in vitro*, but key molecular details, such as how DNA and ubiquitin chains precisely activate SPRTN’s protease function, remain unclear. Overall, DNA is considered to be the main regulator of SPRTN activity^7,8^. SPRTN preferentially recognises specific DNA structures (e.g., ss/dsDNA junctions) to achieve efficient activation^15^. Nevertheless, SPRTN displays relatively low protease activity. *In vitro* cleavage of substrates such as histones, topoisomerases, or model DPC substrates, typically requires hours and often remains incomplete^7,8,15,16^. DNA also induces SPRTN auto-cleavage *in trans*, and auto-cleaved SPRTN is considered to be inactive^8,17^. Knowledge of this on a structural level is limited. The only existing crystal structure (PDB: 6MDX) shows a compact arrangement of the MPD and ZBD domains bound to a small dinucleotide ssDNA with what appears to be limited accessibility to the protease active site^18^. DNA is hypothesised to promote an open conformation of SPRTN, although the detailed mechanism remains largely unknown. Small-angle X-ray scattering (SAXS) has shown that full-length SPRTN becomes more flexible upon DNA-binding, indicating an opening of the protein, but the exact region undergoing allosteric rearrangement remains unclear^8^. The impact of active-site changes on accessibility has not yet been experimentally investigated. Confusingly, in the same study, hydrogen/deuterium exchange mass spectrometry (HDX-MS) determined that the SPRTN C-terminus was more exposed to solvent in the presence of DNA, whereas the protease region was largely protected from hydrogen/deuterium exchange by DNA binding^8^.

The second layer of SPRTN activity regulation involves ubiquitin. In addition to the well-established role of the C-terminal UBZ domain in recruiting ubiquitinated DPC substrates^19,20^, recently, we and others independently identified a new ubiquitin-binding domain in the N-terminal catalytic region, termed the ubiquitin-binding interface domain (USD)^16,21^. The USD binds all types of ubiquitin chains, and this ubiquitin-chain binding acts as a key catalyst that accelerates rapid SPRTN proteolysis following its initial activation by DNA^16,21^. This Ub-chain activation is based on an avidity effect – the longer the Ub chains, the higher the affinity to SPRTN, resulting in the higher activity of SPRTN proteolysis^16^. The USD is located in the MPD domain and, upon binding to Ub chains, directs SPRTN’s proteolytic activity specifically toward ubiquitinated DPCs while sparing key DNA replication components^16^. Molecular dynamics (MD) simulations further suggest that ubiquitin can stabilise an open conformation of the SPRTN protease region^21^. However, a ColabFold-predicted SprT structure comprising the MPD and ZBD domains, which already adopts an open conformation, was used as the starting model for these MD simulations^21^, and the activating effect of DNA was not considered. Because DNA is the primary trigger for SPRTN protease activation, and ubiquitin cannot activate SPRTN in the absence of DNA^16^, it remains unclear how ubiquitin stabilises the open conformation, given that SPRTN is most likely compact without DNA. The most plausible explanation is that the stabilising effect of ubiquitin occurs only after DNA-dependent activation.

For a long time, biochemical characterisation of SPRTN protease activity *in vitro* has been challenging. The recent discoveries of the USD-mediated Ub-activation effect have enabled more detailed investigation of SPRTN protease activity^16,21^, but kinetic data are still largely lacking. Here, based on the identification of a major cleavage site on the histone H1 substrate, we developed a robust peptide-based FRET assay to monitor SPRTN protease kinetics in real time. Combined with other biochemical approaches, we characterised the SPRTN N-terminal protease region (MPD-ZBD-BR) in depth. We found that the auto-cleaved SPRTN N-terminus is highly stable and can still be potentiated by ubiquitin chains. Surprisingly, the isolated MPD domain lost both DNA dependence and Ub avidity, yet retained protease activity. We further discovered that the ZBD domain inhibits the protease activity of the MPD domain, thereby maintaining SPRTN in an autoinhibitory state that cannot be released by DNA alone. Using a site-specific bioorthogonal approach and native mass spectrometry, we provided experimental evidence that DNA, together with the key regulatory elements ZBD and BR, shifts the SPRTN protease region from a closed to an open conformation with a more accessible active site. Together, our work reveals a previously uncharacterised autoinhibitory effect and provides essential missing mechanistic details of how DNA binding induces conformational changes in the SPRTN protease catalytic groove to release the auto-inhibition.

## Results

### Identification of H1 cleavage sites mediated by SPRTN

We recently discovered that SPRTN preferably cleaves the C-terminal tail of H1^16^. To further identify the exact cleavage site, an *in vitro* cleavage assay by a truncated form of SPRTN (SprT-BR; 26-240) (Fig. 1A) was performed using H1 carrying the S191C point mutation as the substrate (Fig. 1B and Supplementary Fig. 1A) with the addition of M1-hexaUb chains as a catalyst^16^. Within 1.5h, the C-terminal H1 can be completely trimmed as a major shorter version of H1 was accumulated (Fig. 1B). Intact mass spectrometry identified several H1 fragments that all end at residue K188, indicating that the major trimmed C-terminal cleavage product by SPRTN proteolysis is “RGCKKK” (aa189-194, Fig. 1C and Supplementary Fig. 1B).

**Figure 1.**
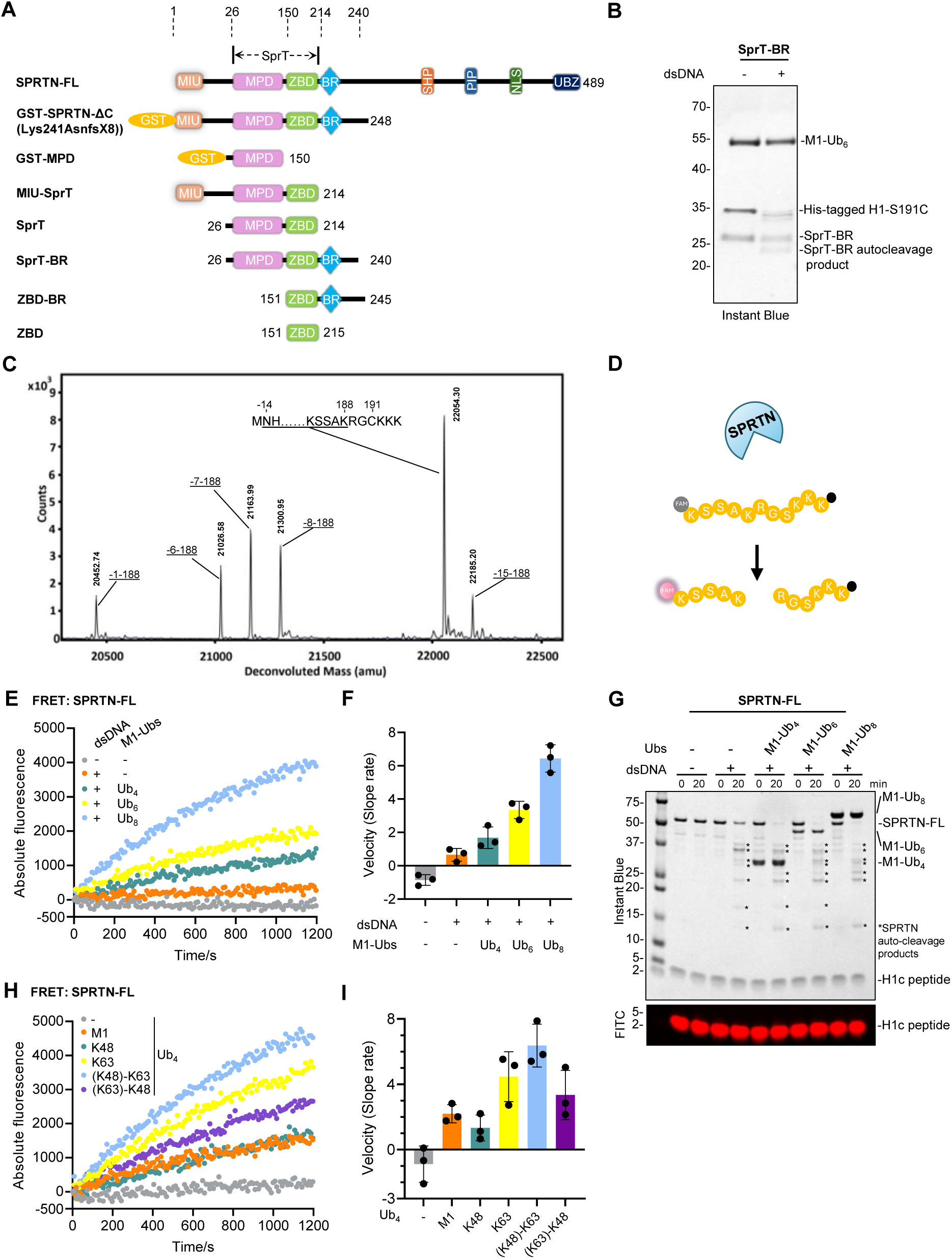
A powerful FRET-based assay reveals real-time kinetics of SPRTN substrate cleavage activated by ubiquitin chains. **(A)** Schematics of SPRTN domain structure and the truncations used for Figure 1-4. MIU, motif Interacting with Ub-binding domain; SprT, the metalloprotease domain similar to that of the *E. coli* SprT protein; MPD, metalloprotease domain; ZBD, zinc binding domain; BR, basic region; PIP, PCNA interaction peptide; SHP, p97 or VCP-binding motif; UBZ, ubiquitin-binding zinc finger. **(B)** SprT-BR cleavage assay towards the model H1 substrate for intact MS. Recombinant SprT-BR (2 µM), His-tagged H1-S191C (1 µM) and M1-hexaUb (2 µM) were incubated in the absence or presence of dsDNA_20/23nt (2.7 µM) at 30°C for 1.5h. The reaction was analysed by SDS-PAGE and Instant Blue staining. **(C)** Intact MS spectrum from the reaction in Figure 1B. The major identified cleavage species are highlighted with corresponding protein boundaries next to them. **(D)** Design of the H1c peptide for the FRET assay. The black dot represents the quencher group Dabcyl. **(E)** Multi-turnover SPRTN cleavage assay towards the H1c peptide monitored by FRET in the presence of M1-linked Ub chains with different lengths. Recombinant full-length SPRTN (2 µM), H1c peptide (20 µM) and M1-linked Ub chains (2 µM) were incubated in the absence or presence of dsDNA_20/23nt (2.7 µM) at 30°C. **(F)** Quantification of the cleavage velocity (slope rate) by linear-fitting the first 4-min curve from Figure 1E. **(G)** SDS-PAGE analysis of the samples from the starting and ending points from the FRET assay performed in Figure 1E. FAM signal from H1c peptide was visualised by FITC scanning on an iBright 1500 imaging system (Invitrogen). **(H)** Multi-turnover SPRTN cleavage assay towards the H1c peptide monitored by FRET in the presence of tetraUb chains with different linkages or branches. Recombinant full-length SPRTN (2 µM), H1c peptide (20 µM) and tetraUb chains (2 µM) were incubated in the presence of dsDNA_20/23nt (2.7 µM) at 30°C. Branched tetraUb: (K48)-K63 indicates [Ub]_2_-^48,63^Ub-^63^Ub; (K63)-K48 indicates [Ub]_2_-^48,63^Ub-^48^Ub. **(I)** Quantification of the cleavage velocity (slope rate) by linear-fitting the first 4-min curve from Figure 1H. The FRET signals for FAM from Figure 1E and 1H were monitored by a platereader (FLUOSTAR, BMG) with the fluorescence mode at Ex. 485 nm and Em. 520 nm for 20 min. Each curve is averaged from 3 repeats. See also Supplementary Figure 1.

### A powerful FRET-based assay reveals real-time kinetics of SPRTN substrate cleavage activated by ubiquitin chains

The existing single-turnover Cy5-based fluorescence assay for SPRTN^16^ (see also Supplementary Fig. 1C) offers improved resolution but remains time-consuming and low-throughput. To address this limitation, we designed a new multi-turnover FRET-based assay using a modified peptide as a model SPRTN substrate, based on the cleaved H1 C-terminal fragment identified by intact mass analysis. Specifically, we selected an 11-amino acid H1 C-terminal tail containing the K188/R189 cleavage site (KSSAK/RGSKKK; hereafter referred to the H1c peptide). The peptide is N-terminally labelled with FAM and C-terminally modified with the quencher Dabcyl. By FRET, the intact peptide exhibits low FAM fluorescence, whereas SPRTN-mediated cleavage separates FAM from Dabcyl, abolishes quenching, and restores full FAM signal (Fig. 1D). Because SPRTN has a relatively weak DNA-induced protease activity, the multi-turnover FRET-based assay (H1c: SPRTN = 20:1) was still not sufficiently sensitive when only DNA was added to the reaction, as indicated by the kinetic data (Figure 1E, orange curve). In contrast, the addition of long ubiquitin chains strongly enhanced SPRTN activity toward the H1c peptide (Figure 1E). This newly developed FRET-based assay thus enables, for the first time, real-time monitoring of multi-turnover SPRTN cleavage kinetics toward its substrate.

Based on this method, the Ub-dependent activation of SPRTN can be quantified and directly compared across conditions (Fig. 1E-F). To determine the multi-turnover rate, the initial velocity (first 4 min) of each kinetic trace in the different Ub-chain contexts was linearly fitted (Fig. 1F). These multi-turnover rates confirmed that ubiquitin-dependent activation of SPRTN follows an avidity manner, consistent with our previous findings^16^. Auto-cleavage of full-length SPRTN was monitored and validated by resolving FRET samples from the start and end points on denaturing SDS–PAGE (Fig. 1G). A conventional single-turnover assay was also performed using H1 C-terminally labelled with Cy5 (H1-C-Cy5, Supplementary Fig. 1A) as a model SPRTN substrate (H1:SPRTN = 1:2). In this assay as well, both substrate cleavage and auto-cleavage confirmed the Ub-dependent activation of SPRTN (Supplementary Fig. 1C). The successful establishment of the FRET assay can be extended to a wider range of applications. For instance, we quantitatively compared the activation effects of tetraUb chains with different linkages or branches, and found that a (K48)-K63 branched tetraUb ([Ub]2-^48,63^Ub-^63^Ub) exhibited the strongest activation of SPRTN (Figure 1H-1I). The activity of truncated SPRTN can also be robustly monitored in the presence of Ub chains (Supplementary Fig. 1D). These results further confirm that DNA is the primary tier of SPRTN activation. Ub chains alone could not activate SprT-BR in the absence of DNA, as demonstrated by both multi-turnover FRET assays (Supplementary Fig. 1D) and single-turnover assays (Supplementary Fig. 1E).

Considering the important role of SPRTN in resolving DPCs, the FRET assay provides a superior platform for inhibitor screening. To validate this concept, a general metalloprotease inhibitor 1,10-phenanthroline was used in an inhibition test. As anticipated, a relatively high concentration (500 µM) of 1,10-phenanthroline inhibited SprT-BR protease activity in the FRET assay (Supplementary Fig. 1F). Interestingly, a low concentration of Zn²⁺ (10 µM) also strongly inhibited SprT-BR activity (Supplementary Fig. 1G). Both effects were confirmed by the single-turnover fluorescence-based assay (Supplementary Fig. 1H). Unlike the intrinsically coordinated Zn²⁺ in SPRTN, free Zn²⁺ likely competes for the catalytic site, as exemplified by carboxypeptidase A (CPA), in which an inhibitory Zn²⁺ forms a complex with the catalytic Zn²⁺ and residue E270^22^.

In summary, we have established a robust FRET-based assay to monitor SPRTN kinetics in real time, enabling detailed characterisation of different SPRTN variants.

### The auto-cleaved SPRTN N-terminal domain remains responsive to ubiquitin chains

It has been previously reported that SPRTN-auto, the major auto-cleaved N-terminal SPRTN product, exhibits protease activity similar to that of SPRTN-ΔC (a patient variant, Lys241AsnfsX8)^8^. However, it remains unclear whether Ub chains can activate SPRTN-auto. Interestingly, the auto-cleaved SPRTN still appeared capable of cleaving the substrate in the presence of Ub chains (Supplementary Fig. 1C: 5-40 min for M1-tetraUb; 5-30 min for M1-hexaUb; 5 min for M1-octaUb). To further investigate this, we performed a FRET assay comparing the activities of full-length and auto-cleaved SPRTN (Fig. 2A). Strikingly, pre-autocleaved SPRTN showed substantial protease activity in the presence of M1-octaUb, only slightly weaker than full-length SPRTN (Fig. 2A). To better visualise both substrate cleavage and SPRTN auto-cleavage, we performed a single-turnover assay. An initial reaction containing all necessary components, including M1-tetraUb, was incubated for one hour to consume nearly all the substrate (H1-C-Cy5) while allowing complete auto-cleavage of SPRTN (Fig. 2B, lane 1-5). After 1 h, a new portion of H1-C-Cy5 with the same amount was added to the reaction. As expected, the substrate was processed again over time, albeit with slightly slower kinetics (Fig. 2B, lane 6-10; Fig. 2C).

**Figure 2.**
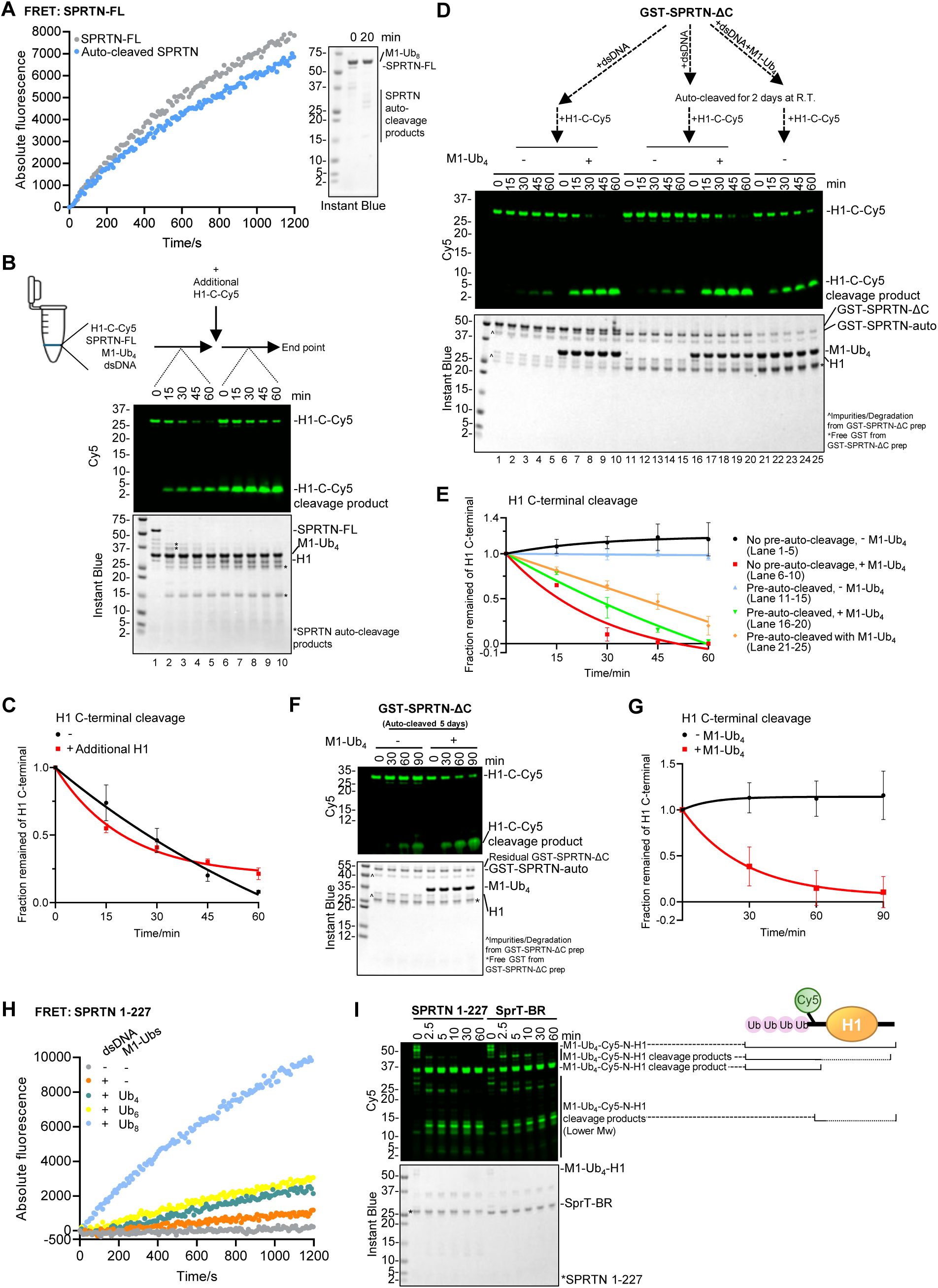
The auto-cleaved SPRTN N-terminal domain remains respsonsive to ubiquitin chains. **(A)** Multi-turnover SPRTN cleavage assay towards the H1c peptide monitored by FRET. Recombinant full-length or auto-cleaved SPRTN (2 µM), H1c peptide (20 µM), M1-octaUb (2 µM) and dsDNA_20/23nt (2.7 µM) were incubated at 30°C. Right panel: Autocleaved SPRTN was made by incubating recombinant full-length SPRTN (2 µM), M1-octaUb (2 µM) and dsDNA_20/23nt (2.7 µM) at R.T. for 20 min. **(B)** Single-turnover SPRTN cleavage assay towards H1-C-Cy5. Recombinant full-length SPRTN (2 µM), H1-C-Cy5 (1 µM), M1-tetraUb (2 µM) and dsDNA_20/23nt (2.7 µM) were incubated for the indicated time at 30°C. By the end of the reaction (1h), an additional same amount of H1-C-Cy5 was added and incubated for another hour with the same condition. Representative figure from 3 repeats. **(C)** Cleavage kinetics of the full-length H1-C-Cy5 substrate (C-terminal cleavage rate) from Figure 2B. n=3. Error bar, SD. **(D)** Single-turnover GST-SPRTN-ΔC cleavage assay towards H1-C-Cy5 with or without pre-auto-cleavage. Lane 1-10: Recombinant GST-SPRTN-ΔC (2 µM), H1-C-Cy5 (1 µM) and dsDNA_20/23nt (2.7 µM) were incubated in the absence or presence of M1-tetraUb (2 µM) for the indicated time at 30°C. Lane 11-20: Recombinant GST-SPRTN-ΔC (2 µM) was pre-auto-cleaved with dsDNA_20/23nt (2.7 µM) at R.T. for 2 days and then added with H1-C-Cy5 (1 µM) for substrate cleavage in the absence or presence of M1-tetraUb (2 µM) for the indicated time at 30°C. Lane 21-25: Recombinant GST-SPRTN-ΔC (2 µM) was pre-auto-cleaved with dsDNA_20/23nt (2.7 µM) in the presence of M1-tetraUb (2 µM) at R.T. for 2 days and then added with H1-C-Cy5 (1 µM) for the indicated time at 30°C. Representative figure from 3 repeats. **(E)** Cleavage kinetics of the full-length H1-C-Cy5 substrate (C-terminal cleavage rate) from Figure 2D. n=3. Error bar, SD. **(F)** Single-turnover GST-SPRTN-ΔC cleavage assay towards H1-C-Cy5 with 5-day pre-auto-cleavage. Recombinant GST-SPRTN-ΔC (2 µM) was pre-auto-cleaved with dsDNA_20/23nt (2.7 µM) at R.T. for 5 days and then added with H1-C-Cy5 (1 µM) in the absence or presence of M1-tetraUb (2 µM) for the indicated time at 30°C. Representative figure from 3 repeats. **(G)** Cleavage kinetics of the full-length H1-C-Cy5 substrate (C-terminal cleavage rate) from Figure 2F. n=3. Error bar, SD. **(H)** Multi-turnover SPRTN 1-227 cleavage assay towards the H1c peptide monitored by FRET. Recombinant SPRTN 1-227 (2 µM), H1c peptide (20 µM) and M1-linked Ub chains (2 µM) were incubated in the absence or presence of dsDNA_20/23nt (2.7 µM) at 30°C. **(I)** Singe-turnover SPRTN cleavage assay towards the model modified substrate M1-Ub_4_-Cy5-N-H1. Recombinant SPRTN 1-227 or SprT-BR (2 µM), M1-Ub_4_-Cy5-N-H1 (1 µM) and dsDNA_20/23nt (2.7 µM) were incubated for the indicated time at 30°C. Representative figure from 3 repeats. The FRET signals for FAM from Figure 2A and 2H were monitored by a platereader (POLARSTAR, BMG) with the fluorescence mode at Ex. 485 nm and Em. 520 nm) for 20 min. Each curve is averaged from 3 repeats. The reactions from Figure 2B, 2D, 2F and 2I were analysed with SDS-PAGE followed by Cy5-scanning on Typhoon FLA 9500 (GE Healthcare) and Instant Blue staining. The Cy5 signals from Figure 2B, 2D, and 2F were analysed by ImageJ. Kinetic data were fitted with one phase exponential decay - least squares fit (Prism). n=3. Error bar, SD. See also Supplementary Figure 2.

To better characterise SPRTN-auto, we first tested the auto-cleavage of GST-SPRTN-ΔC (Fig. 1A). We confirmed that GST-SPRTN-ΔC could generate the same SPRTN-auto species in the presence of dsDNA (Supplementary Fig. 2A, B). Adding M1-tetraUb accelerated both the auto-cleavage and substrate cleavage activity from GST-SPRTN-ΔC (Fig. 2D, compare lanes 1-5 with lanes 6-10; Supplementary Fig. 2A, B).

To isolate SPRTN-auto, GST-SPRTN-ΔC was allowed to auto-cleave with a longer time course. After 24 h at room temperature, less than 25% of GST-SPRTN-ΔC remained (Supplementary Fig. 2A, B). Nevertheless, this mixture of GST-SPRTN-ΔC and SPRTN-auto could still be activated by M1-tetraUb, with only a slightly reduced activity compared with GST-SPRTN-ΔC alone (Supplementary Fig. 2C, lanes 6-10 and 16-20; Supplementary Fig. 2D), suggesting that SPRTN-auto can also be potentiated by Ub chains.

To achieve near-complete conversion to SPRTN-auto, GST-SPRTN-ΔC was incubated with DNA for 2 days at room temperature, either in the absence or presence of M1-tetraUb. Without M1-tetraUb, the substrate-cleaving activity of SPRTN-auto alone was as weak as that of GST-SPRTN-ΔC (Fig. 2D, lanes 1-5 and 11-15; Fig. 2E), consistent with a previous study^8^. In contrast, the protease activity of SPRTN-auto was stimulated by M1-tetraUb (Fig. 2D, lanes 16-20). Remarkably, even after 5 days of auto-cleavage at room temperature, SPRTN-auto still retained Ub-dependent activation (Fig. 2F, G).

Short incubation with M1-tetraUb did not substantially impair SPRTN-auto, whereas prolonged incubation with Ub chains promoted further auto-cleavage of SPRTN-auto (Supplementary Fig. 2A, B). Consequently, the reduced amount of SPRTN-auto led to weaker substrate-cleaving activity (Fig. 2D, lanes 21-25; Fig. 2E). Together, these data indicate that the auto-cleaved N-terminus of SPRTN, which contains the protease domain, is in fact highly stable. SPRTN-auto is resistant to further auto-cleavage unless Ub chains are present (Supplementary Fig. 2A, B).

The major auto-cleavage site in the N-terminus of SPRTN has previously been mapped between K227 and L228^7,18^. To this end, we also assessed the Ub-dependent activation of SPRTN 1-227, a mimic of SPRTN-auto. Although the intrinsic protease activity of SPRTN 1-227 was relatively weak, it retained a strong capacity for activation by Ub chains in both multi-turnover and single-turnover assays (Fig. 2H and Supplementary Fig. 2E, F). The rapid cleavage kinetics of SPRTN 1-227 toward the Ub-modified model substrate M1-Ub4-Cy5-N-H1 (Supplementary Fig. 2G) were also comparable to those of SprT-BR (Fig. 2I).

Together, these detailed characterisations indicate that the auto-cleaved SPRTN N-terminus remains stable, in contrast to the widely accepted dogma of its ‘inactivation’^8,17^. Moreover, the Ub-dependent activation potential of SPRTN-auto remains highly selective for ubiquitinated substrates.

### The isolated MPD protease domain is independent of both DNA activation and Ub avidity effects

It is puzzling that the high stability of SPRTN-auto contrasts with the instability of the MPD domain in isolation, which was previously reported to be difficult to express and purify^18^. Crystals could only be obtained from a longer SPRTN construct, SprT (26-214)^18^. Despite the difficulty in isolating MPD, we successfully purified a GST-tagged MPD domain with improved stability, which enabled us to characterise its protease activity as well as the DNA- and Ub-binding properties in much greater detail.

As expected, MPD, as a typical protease domain, exhibited a basal level of intrinsic activity toward the model substrate H1 (Fig. 3A, lanes 1-8). Cleavage of H1 was not restricted to the extreme C-terminal tail but extended into longer C-terminal regions, as indicated by multiple Cy5-visible cleavage products (Fig. 3A, indicated by white dots). Weak cleavage at the H1 N-terminus using Cy5-N-H1 (Supplementary Fig. 2G) was also observed at very late time points (Supplementary Fig. 3F, lanes 1-7). Unexpectedly, the protease activity of the MPD was no longer DNA-dependent (Fig. 3A, lanes 1-8). Instead, DNA, including both ssDNA and dsDNA, inhibited MPD activity in a dose-dependent manner (Fig. 3A, lanes 9-16; Supplementary Fig. 3A). The DNA-independent activity of the MPD resembled the ssDNA-activated SprT-BR cleavage pattern (Supplementary Fig. 3B), suggesting that ssDNA may induce a conformation of the SPRTN protease region similar to that of the isolated MPD. As expected, ZnCl₂ could still inhibit the protease activity at a low concentration (Supplementary Fig. 3C).

**Figure 3.**
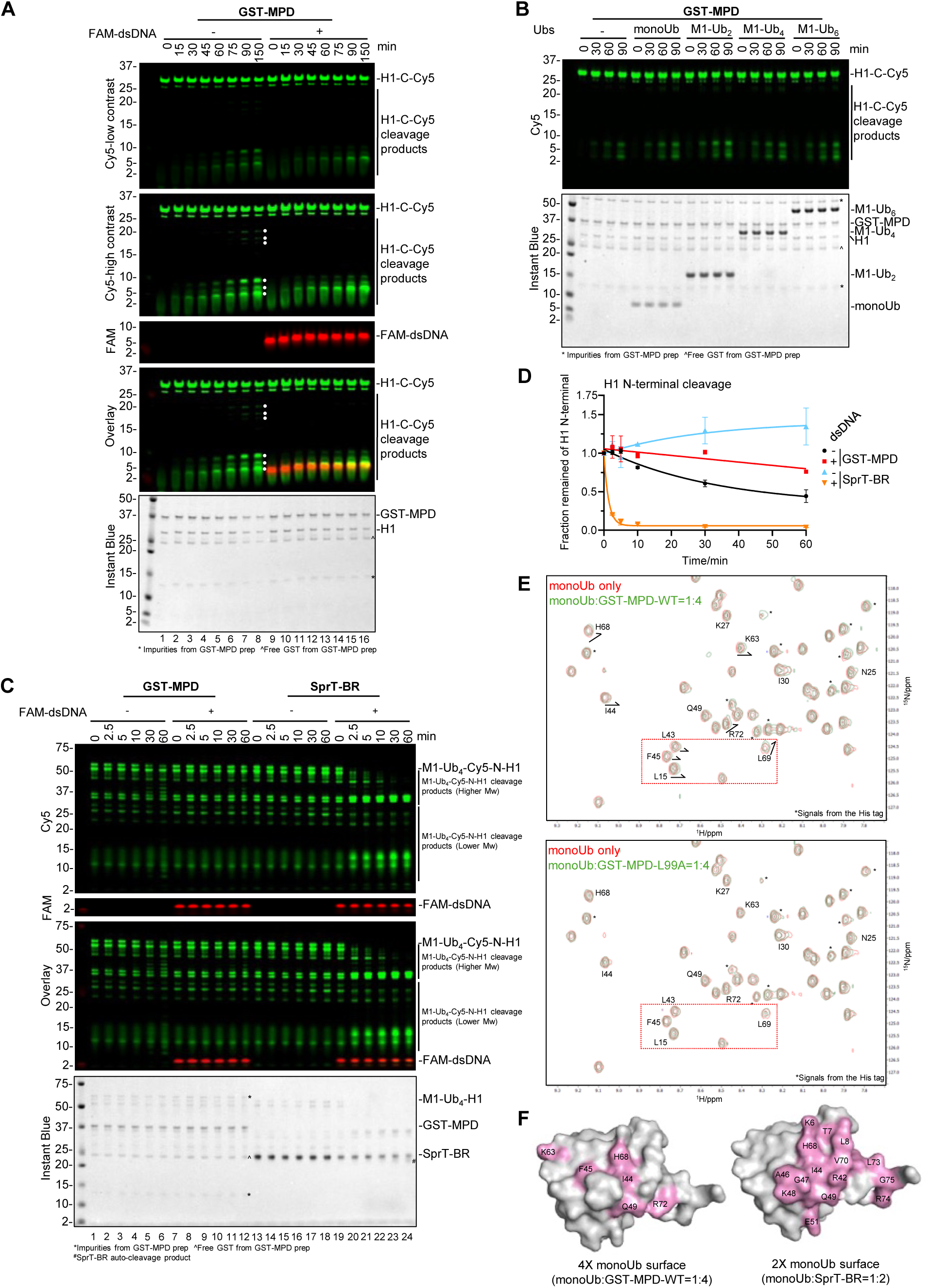
The isolated MPD protease domain is independent of both DNA activation and Ub avidity effects. **(A)** Single-turnover GST-MPD cleavage assay towards H1-C-Cy5. Recombinant GST-MPD (2 µM) and H1-C-Cy5 (1 µM) were incubated in the absence or presence of FAM-labelled dsDNA_20/23nt (2.7 µM) for the indicated time at 30°C. Cleavage products are indicated with white dots. Representative figure from 3 repeats. **(B)** Single-turnover GST-MPD cleavage assay towards H1-C-Cy5 with Ubs. Recombinant GST-MPD (2 µM), H1-C-Cy5 (1 µM) and M1-linked Ubs (2 µM) were incubated for the indicated time at 30°C. DNA is not added. The reactions were analysed by SDS-PAGE, followed by Cy5 scanning on Typhoon FLA 9500 (GE Healthcare) and Instant Blue staining. Representative figure from 3 repeats. **(C)** Single-turnover SPRTN cleavage assay towards the model modified substrate M1-Ub_4_-Cy5-N-H1. M1-Ub_4_-Cy5-N-H1 (1 µM) were incubated with recombinant GST-MPD or SprT-BR (2 µM) in the absence or presence of FAM-labelled dsDNA_20/23nt (2.7 µM) for the indicated time at 30°C. Representative figure from 3 repeats. **(D)** Cleavage kinetics of the full-length M1-Ub_4_-Cy5-N-H1 substrate (N-terminal cleavage rate) from Figure 3C. Cy5 signals from the full-length M1-Ub_4_-Cy5-N-H1 were analysed by the iBright Analysis Software (Invitrogen). Kinetic data were fitted with one phase exponential decay - least squares fit (Prism). N=3. Error bar, SD. **(E)** Superimposition of the 950 MHz ^1^H-^15^N HSQC spectra of monoUb alone (in red contours), monoUb with the addition of GST-MPD-WT (upper) or GST-MPD-L99A (bottom) at a molar ratio of 1:4 (in green contours). Spectra were plotted by MestReNova. The signals were assigned according to the reported chemical shifts of monoUb (BMRB entry: 17769). *Signals from the N-terminal His tag on monoUb. **(F)** Comparison of the perturbed monoUb surface by GST-MPD-WT and SprT-BR. Left: residues with visible chemical shifts upon the addition of GST-MPD at a molar ratio of 1:4 from Figure 3E are highlighted in pink on the monoUb surface (PDB: 1UBQ). Right: residues with complete broadening upon the addition of SprT-BR at a molar ratio of 1:2 are highlighted in pink on the monoUb surface (data recreated from Song et al., 2025). The reactions from Figures 3A and 3C were analysed by SDS-PAGE, followed by Cy5 and FAM scanning on Typhoon FLA 9500 (GE Healthcare) and Instant Blue staining. See also Supplementary Figure 3.

In our previous study, we characterised the interaction between SprT-BR (26-240) and Ub, and identified residue L99, located within the MPD domain, as one of the key residues involved in this interaction^16^. Here, we further examined the effect of Ub on the isolated MPD. Surprisingly, Ub-mediated activation of the MPD was very limited. Only slight activation by Ub chains was observed at late time points, based on quantification of total H1 cleavage products (Fig. 3B and Supplementary Fig. 3D). Notably, the MPD lost the Ub avidity effect, as Ub chains of different lengths had minimal impact (Fig. 3B and Supplementary Fig. 3D), indicating that the MPD likely needs to be properly positioned by neighbouring domains or factors to achieve maximal Ub avidity. This is consistent with a recent NMR study showing that residue I212 in the ZBD domain undergoes perturbation upon addition of monoUb to SprT-BR^21^. The effect of Ub was also dose-independent (Supplementary Fig. 3E).

We also tested the activity of the MPD toward the Ub-modified model substrate M1-Ub4-Cy5-N-H1. In contrast to the rapid substrate cleavage by SprT-BR in the presence of DNA, the MPD displayed much slower kinetics, even in the absence of DNA (Fig. 3C, compare lanes 1-6 with 19-24; Fig. 3D; Supplementary Fig. 3F, lanes 8-14), consistent with the results in Fig. 3B. Again, the protease activity of the MPD was inhibited by DNA (Fig. 3C, compare lanes 1-6 with 7-12; Fig. 3D).

All of these observations raised our concern that the interaction between Ub and the MPD is likely to be very weak. To further characterise this interaction and validate the binding surface on Ub, we performed NMR titration experiments in which GST-MPD-WT (or the L99A variant, or free GST) was titrated into ^15^N-labelled monoUb (Fig. 3E and Supplementary Fig. 3G). From the 2D-HSQC spectra, only a few residues on the monoUb surface showed weak chemical shift changes upon addition of excess GST-MPD-WT, whereas the GST-MPD-L99A variant did not induce any detectable perturbation in monoUb (Fig. 3E). The residues exhibiting perturbation upon 4-fold addition of GST-MPD-WT binding were mapped onto the monoUb surface and compared with the perturbed residues upon 2-fold addition of SprT-BR (data recreated from Song et al., 2025). The perturbed residues on monoUb in the presence of GST-MPD-WT were very limited (Fig. 3F), confirming that the binding between monoUb and MPD is indeed very weak.

In summary, the isolated MPD domain, which possesses basal intrinsic protease activity, functions independently of both DNA-mediated activation and Ub-avidity effect. These results highlight the importance of additional N-terminal modules of SPRTN in regulating its protease activity.

### The SprT domain is catalytically inactive but can be weakly reactivated by longer ubiquitin chains

We next characterised the SPRTN N-terminal region lacking the BR motif in more detail. It was previously reported that MIU-SprT (1-214) is inactive^18^, suggesting that BR plays an essential role in activating the DNA-dependent protease activity^15,18^. In our conventional fluorescence assay, MIU-SprT displayed almost undetectable intrinsic protease activity, even in the presence of dsDNA (Supplementary Fig. 4A). However, the addition of longer Ub chains weakly stimulated protease activity, indicating that MIU-SprT still exhibits the Ub-avidity effect (Supplementary Fig. 4A). The shortest SprT domain (26-214) behaved similarly (Supplementary Fig. 4B). Although MD simulations have suggested that Ub tends to stabilise the open conformation of SprT^21^, the Ub-dependent activation of the SprT domain observed here was very weak, despite the Ub-binding region from the MPD domain is still present. This suggests that the SprT domain does not adopt a fully open conformation, even in the presence of DNA.

### The SprT domain exerts an autoinhibitory effect

Considering that the isolated MPD domain represents the core protease domain for SPRTN activity (Fig. 3A; Fig. 4A, lanes 1-8), the suppressed activity of SprT suggests an intramolecular inhibitory effect from the ZBD on the MPD (Fig. 4A, lanes 9-16; Supplementary Fig. 4B). This auto-inhibition was also observed in the SprT-BR construct in the absence of DNA (Fig. 4A, lanes 17-20). In the presence of DNA, SprT alone failed to overcome this inhibition, whereas SprT-BR regained substantial activity, indicating that DNA binding to the BR domain is critical to activate SPRTN (Fig. 4A, lanes 17-24). Altogether, these results imply that DNA, together with the DNA-binding modules MPD and BR, plays an essential role in regulating SPRTN protease activity.

**Figure 4.**
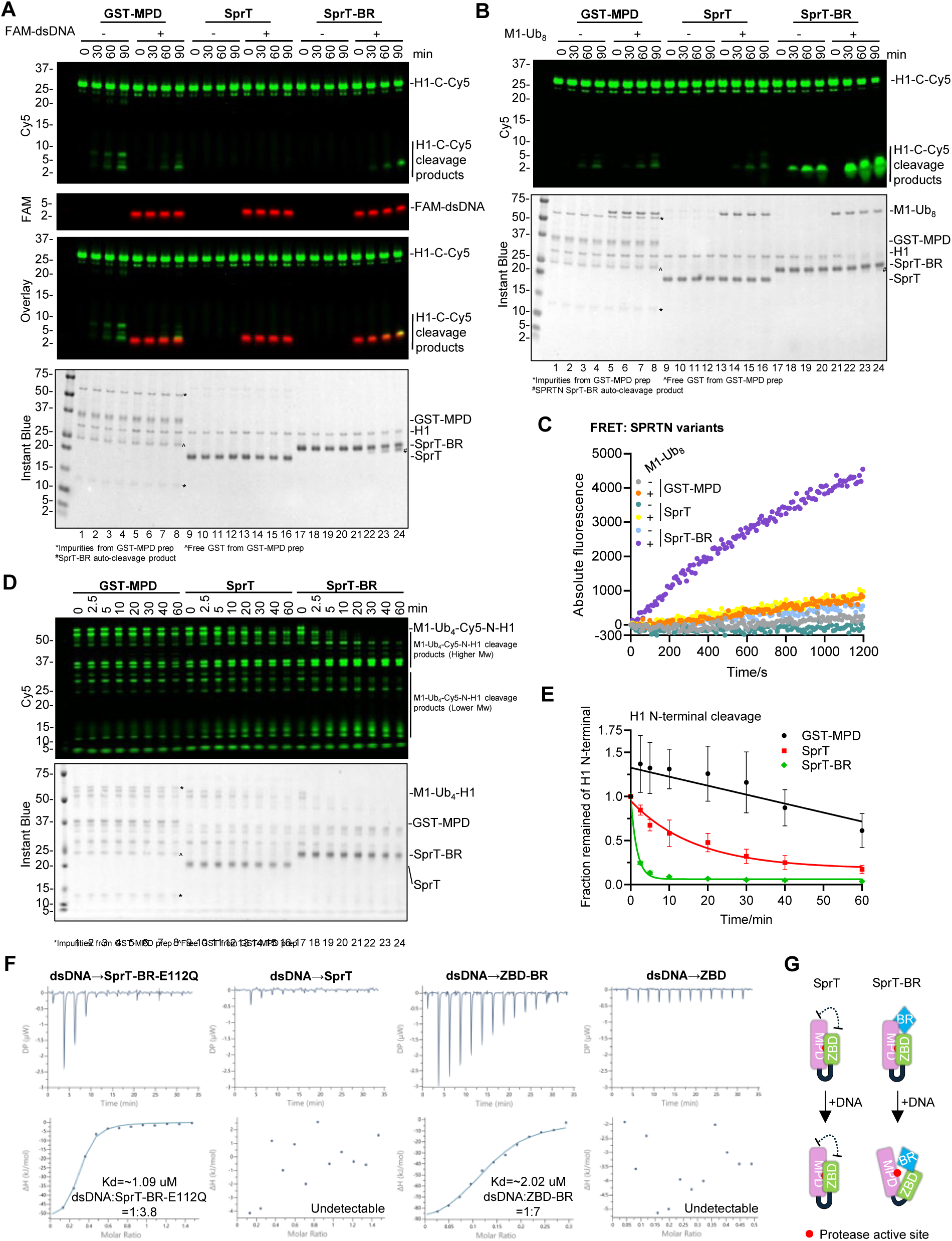
The steric effect between ZBD and MPD maintains the SPRTN N-terminal protease region in auto-inhibition. **(A)** Activity comparison by single-turnover SPRTN cleavage assay towards H1-C-Cy5. Recombinant SPRTN variants (GST-MPD, SprT or SprT-BR, 2 µM) and H1-C-Cy5 (1 µM) were incubated in the absence or presence of FAM-labelled dsDNA_20/23nt (2.7 µM) for the indicated time at 30°C. Representative figure from 3 repeats. **(B)** Comparison of Ub-activation effect by single-turnover SPRTN cleavage assay towards H1-C-Cy5. Recombinant SPRTN variants (GST-MPD, SprT or SprT-BR, 2 µM), H1-C-Cy5 (1 µM) and dsDNA_20/23nt (2.7 µM) were incubated in the absence or presence of M1-octaUb (2 µM) for the indicated time at 30°C. Representative figure from 3 repeats. **(C)** Multi-turnover SPRTN cleavage assay towards the H1c peptide monitored by FRET. Recombinant SPRTN variants (GST-MPD, SprT or SprT-BR, 2 µM), H1c peptide (20 µM) and dsDNA_20/23nt (2.7 µM) were incubated in the absence or presence of M1-octaUb (2 µM) at 30°C. The FRET signals for FAM were monitored by a platereader (POLARSTAR, BMG) with the fluorescence mode at Ex. 485 nm and Em. 520 nm for 20 min. Each curve is averaged from 3 repeats. **(D)** Single-turnover SPRTN cleavage assay towards the model modified substrate M1-Ub_4_-Cy5-N-H1. Recombinant SPRTN variants (GST-MPD, SprT or SprT-BR, 2 µM) and M1-Ub_4_-Cy5-N-H1 (1 µM) were incubated in the presence of dsDNA_20/23nt (2.7 µM) for the indicated time at 30°C. Representative figure from 3 repeats. **(E)** Cleavage kinetics of the full-length M1-Ub_4_-Cy5-N-H1 substrate (N-terminal cleavage rate) from Figure 4D. Cy5 signals from the full-length M1-Ub_4_-Cy5-N-H1 were analysed by the iBright Analysis Software (Invitrogen). Kinetic data were fitted with one phase exponential decay - least squares fit (Prism). N=3. Error bar, SD. **(F)** Isothermal titration calorimetry (ITC) analysis of the interaction between dsDNA_20/23nt and different SPRTN N-terminal variants. The dissociation constant (Kd) and stoichiometry of binding (N) are indicated here and summarised in Table S1A. **(G)** Schematic of the model showing the auto-inhibition within SprT (left) and the model of the possible involvement of BR in the release of auto-inhibition (right).= The reactions from Figure 4A-B and 4D were analysed by SDS-PAGE, followed by Cy5 scanning on Typhoon FLA 9500 (GE Healthcare) and Instant Blue staining. SDS-PAGE from Figure 4A was additionally analysed by FAM scanning on Typhoon FLA 9500 (GE Healthcare). See also Supplementary Figure 4.

We also compared the Ub-dependent activation of SprT and SprT-BR. As expected, SprT could not be effectively activated by M1-octaUb, whereas the activity of SprT-BR was significantly stimulated (Figure 4B, compare lanes 13-16 with 21-24). A FRET assay confirmed these findings, revealing only minor activation of MPD activity by M1-octaUb (Fig. 4C). Using M1-Ub4-fused H1 as a model ubiquitinated substrate (Supplementary Fig. 2G), we observed that SprT exhibited much slower substrate-cleavage kinetics than SprT-BR (Fig. 4D, E, compare lanes 9-16 with 17-24). Several high-molecular-weight cleavage products accumulated but were not further processed by SprT within 1 h (Fig. 4D, lanes 9-16). Collectively, these observations indicate that the SprT domain remains largely inhibited, whereas DNA relieves the auto-inhibition of SprT-BR, possibly by promoting a more open conformation. In addition, Ub chains may exert a stronger activating effect by stabilising SprT-BR in this open conformation.

### MPD and BR redefine the DNA-binding stoichiometry of ZBD

Comparison of the NMR spectra of SprT-BR and ZBD-BR highlights key residues with large perturbations on the ZBD surface^21^. The crystal structure of SprT in complex with a 2-nt ssDNA (PDB: 6MDX) also reveals a compact fold between MPD and ZBD stabilised by extensive hydrophobic interactions^18^. Collectively, these data imply a steric effect between MPD and ZBD within the SprT construct, which appears to be maintained in a relatively closed conformation. However, titration with either isolated

ZBD or ZBD-BR did not inhibit MPD activity at a 1:1 ratio (Supplementary Fig. 4C), suggesting that a direct intramolecular interaction between ZBD and MPD is not strictly required. Instead, the steric effect is most likely mediated by the flexible linker, which constrains the two subdomains within a limited space.

Intriguingly, the DNA-binding ability of MIU-SprT is undetectable by EMSA^18^. NMR can detect weak binding between ssDNA and SprT, but the perturbations are minimal compared with the robust interaction observed between ssDNA and SprT-BR^18^. In contrast, binding between ssDNA and isolated ZBD is readily detectable by NMR^15^. This is puzzling, given that ZBD is the major DNA-binding module in SPRTN. Therefore, it underscores the need to quantitatively compare the affinity and stoichiometry of DNA binding across different SPRTN constructs.

We employed ITC to measure the DNA-binding affinities of SprT-BR, SprT, ZBD-BR, and ZBD respectively. Consistent with the lack of detectable binding by EMSA^18^, the interaction between DNA and SprT was too weak to be observed by ITC (Fig. 4F and Supplementary Fig. 4D). In contrast, a 20-nt ssDNA bound the isolated ZBD with a Kd of 0.84 µM and an apparent binding ratio of approximately 1:19 (Supplementary Fig. 4D and Table S1A), whereas ZBD did not bind dsDNA (Fig. 4F). This is consistent with the SprT crystal structure (PDB: 6MDX), in which the ZBD is positioned to interact with ssDNA bases, most likely one ZBD molecule per one base. These observations indicate that, within the SprT construct, the MPD largely restrains the DNA-binding from the ZBD, probably due to the overall bulk and compactness of SprT. ZBD-BR also displayed appreciable affinity to DNA, but with a reduced binding ratio of 7:1 for dsDNA (Fig. 4F and Table S1A) and 1.9:1 for ssDNA (Supplementary Fig. 4D and Table S1A), suggesting that BR also contributes to DNA-binding stoichiometry. SprT-BR-E112Q bound both ssDNA and dsDNA with moderate affinity (∼1-4 μM) and a binding stoichiometry of ∼2:1-4:1^16,18^ (Fig. 4F and Table S1A). The noticeable differences in stoichiometry across constructs suggest that the MPD and BR together remodel SPRTN to engage defined amounts of DNA. This likely enables SPRTN to act in proximity to DNA economically, consistent with its low abundance *in vivo*.

In summary, the steric interplay between MPD and ZBD maintains the SPRTN protease region in a tightly controlled state. ZBD suppresses the intrinsic activity of MPD, whereas MPD in turn limits the DNA-binding capacity of ZBD. BR plays an essential role in relieving this auto-inhibition in the presence of DNA. ZBD and BR confer the DNA-dependent properties of SPRTN, while MPD and BR jointly redefine its DNA-binding stoichiometry (Fig. 4G).

### The ZBD-BR-DNA trinity relieves auto-inhibition *in cis*

We further investigated the mechanism by which BR relieves auto-inhibition. Compared with SprT, the DNA-activated protease activity of SprT-BR was higher (Fig. 4A). We therefore first examined the DNA-binding affinity of a synthesised BR peptide. Surprisingly, the affinity between the BR peptide and DNA was extremely low, with a Kd of >15 mM by MST (Supplementary Fig. 5A and Table S1B). BR’s affinity for dsDNA was much lower than for ssDNA, consistent with previous NMR observations^15^.

Comparison of 2D-HSQC NMR spectra of ZBD and ZBD-BR also suggests that BR has transient interactions with ZBD^15^. MST measurements confirmed weak binding between the isolated ZBD and the BR peptide, with a Kd of ∼2 mM (Fig. 5B and Table S1B). It is possible that the full-context dependent effect of the BR can not be replicated using an isolated peptide. Thus, to further validate the importance of the ZBD-BR interaction, we generated a chimeric BR-SprT construct in which BR was fused to the N-terminus of SprT, thereby mimicking SprT-BR (Fig. 5A). In the presence of dsDNA, BR-SprT partially rescued the weak activity of SprT, but its activity remained lower than the basal level of the isolated MPD in the absence of DNA (Fig. 5C and Supplementary Fig. 5B). In the presence of long Ub chains, both the Ub-dependent activation and Ub avidity effects were weakly restored in BR-SprT (Fig 5D and Supplementary Fig. 5C, compared with Supplementary Fig. 4A, B). The cleavage kinetics of BR-SprT toward M1-Ub4-Cy5-N-H1 were also partially recovered (Fig. 5E, F). Collectively, these results indicate that BR, ZBD, and DNA act together to induce conformational rearrangements in the SPRTN protease region, likely exposing the catalytic groove to facilitate substrate cleavage.

**Figure 5.**
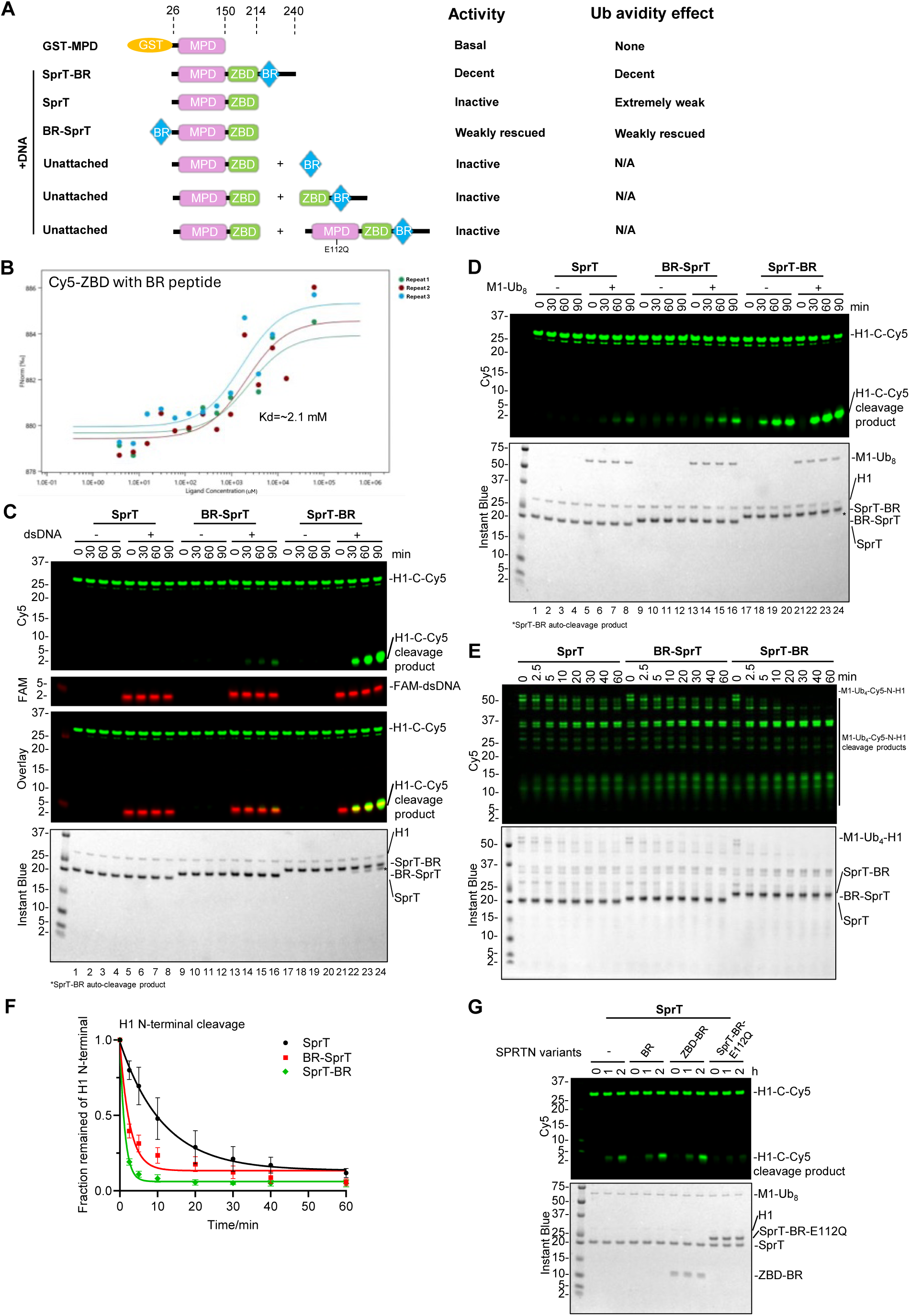
The ZBD-BR-DNA trinity relieves auto-inhibition *in cis*. **(A)** Schematic of the SPRTN constructs and experiment design. The protease activity and Ub avidity effect are also summarised. **(B)** MST affinity analysis between BR peptide and Cy5-ZBD under 20% LED power. The dissociation constant (Kd) is indicated here and summarised in Table S1B. **(C)** Activity comparison by single-turnover SPRTN cleavage assay towards H1-C-Cy5. Recombinant SPRTN variants (SprT, BR-SprT or SprT-BR, 2 µM) and H1-C-Cy5 (1 µM) were incubated in the absence or presence of FAM-labelled dsDNA_20/23nt (2.7 µM) for the indicated time at 30°C. Representative figure from 3 repeats. **(D)** Comparison of Ub-activation effect by single-turnover SPRTN cleavage assay towards H1-C-Cy5. Recombinant SPRTN variants (SprT, BR-SprT or SprT-BR, 2 µM), H1-C-Cy5 (1 µM) and dsDNA_20/23nt (2.7 µM) were incubated in the absence or presence of M1-octaUb (2 µM) for the indicated time at 30°C. Representative figure from 3 repeats. **(E)** Single-turnover SPRTN cleavage assay towards the model modified substrate M1-Ub_4_-Cy5-N-H1. Recombinant SPRTN variants (SprT, BR-SprT or SprT-BR, 2 µM) and M1-Ub_4_-Cy5-N-H1 (1 µM) were incubated in the presence of dsDNA_20/23nt (2.7 µM) for the indicated time at 30°C. Representative figure from 3 repeats. **(F)** Cleavage kinetics of the full-length M1-Ub_4_-Cy5-N-H1 substrate (N-terminal cleavage rate) from Figure 5E. Cy5 signals from the full-length M1-Ub_4_-Cy5-N-H1 were quantified by the iBright Analysis Software (Invitrogen). Kinetic data were fitted with one phase exponential decay - least squares fit (Prism). N=3. Error bar, SD. **(G)** Single-turnover SprT cleavage assay towards H1-C-Cy5 in the presence of BR, ZBD-BR or SprT-BR-E112Q. Recombinant SprT (2 µM), H1-C-Cy5 (1 µM), M1-octaUb (2 µM) and dsDNA_20/23nt (2.7 µM) were incubated with different SPRTN variants (BR peptide, ZBD-BR or SprT-BR-E112Q, 2 µM) for the indicated time at 30°C. Representative figure from 3 repeats. The reactions from Figure 5C-E and 5G were analysed by SDS-PAGE, followed by Cy5 scanning on Typhoon FLA 9500 (GE Healthcare) and Instant Blue staining. SDS-PAGE from Figure 5C was additionally analysed by FAM scanning on Typhoon FLA 9500 (GE Healthcare). See also Supplementary Figure 5.

To determine whether the ZBD-BR-DNA trinity functions *in cis* or *in trans*, BR, ZBD-BR, or SprT-BR-E112Q was titrated into SprT at a 1:1 ratio in the presence of DNA (Fig. 5A, G). None of these three SPRTN constructs was able to activate SprT (Fig. 5G), indicating that ZBD and BR function effectively only in an intramolecular (*cis*) context.

Collectively, BR is the key element that relieves auto-inhibition through two essential mechanisms: (i) binding to DNA and (ii) intramolecular proximity to ZBD. In this way, the ZBD-BR-DNA trinity tightly regulates the protease activity of SPRTN *in cis*.

### The ZBD-BR-DNA trinity induces an open conformation that relieves auto-inhibition

Over the past few years, solution NMR has been widely used to study the dynamics of SPRTN, particularly its DNA-binding properties. Although residue assignments for the ZBD and BR domains are available^15,21^, resolving the SprT-BR structure by NMR remains challenging due to the pronounced instability of MPD. Without complete residue assignments and additional constraints from more complex 3D NMR spectra, it is difficult to provide direct evidence of DNA-induced conformational changes, particularly the opening of the structure.

To gain further insight into the DNA-binding dynamics of the SPRTN N-terminus, native MS was employed to analyse the distribution of solvent-accessible surface area for SPRTN variants in complex with dsDNA (Fig. 6A, B and Supplementary Fig. 6A, B). Binding of dsDNA to SprT-BR-E112Q resulted in a major mass-to-charge shift and, more importantly, a broader surface area distribution, indicating a more exposed, flexible structure (Fig. 6B). In contrast, only a small fraction of SprT exhibited a mass-to-charge shift upon binding to dsDNA, and this population showed more flexibility (Supplementary Fig. 6A, B), consistent with its weak interaction with DNA.

**Figure 6.**
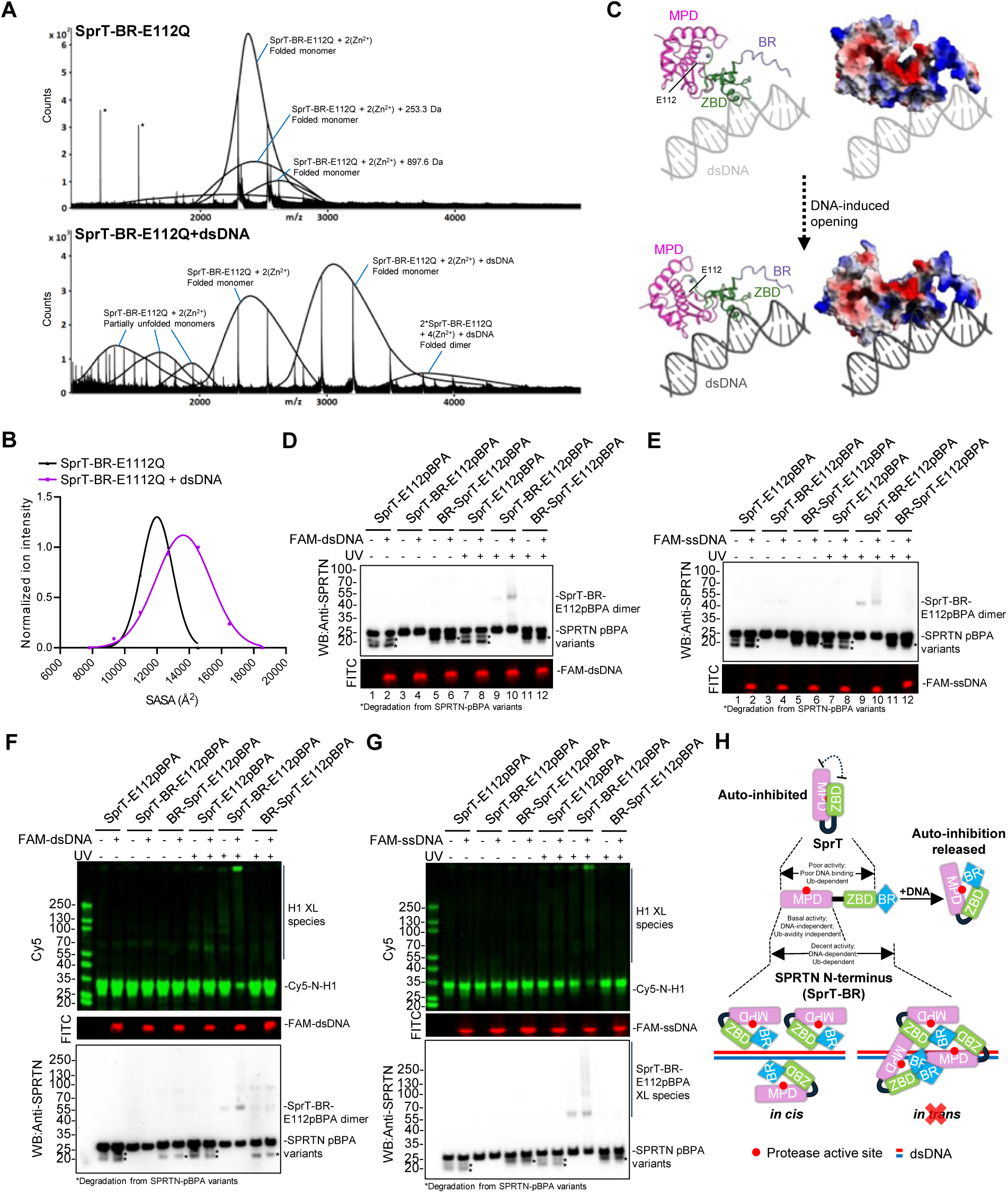
The ZBD-BR-DNA trinity induces an open conformation of SPRTN-BR that relieves auto-inhibition. **(A)** Native MS analysis of SprT-BR-E112Q (upper) or in complex with dsDNA_20/23nt (lower). *Tune mix standards. **(B)** Gaussian distribution of the solvent accessible surface area (SASA^2^) from Figure 6A. Data were fitted by Caussian – least squares fit (Prism). **(C)** Modelling of an open conformation of SprT-BR induced by dsDNA in cartoon mode (Left) and surface mode (Right). The charged surface is colour-coded by red-blue spectrum (Red: negatively charged; Blue: positively charged). The upper starting model was built based on extending dsDNA from the SprT-DNA crystal structure (PDB: 6MDX). Please see Supplementary Figure 6D. **(D)** SPRTN-pBPA trapping assay by dsDNA. SPRTN variants (∼ 0.18 µM) carrying the E112pBPA modification were incubated in the absence or presence of FAM-labelled dsDNA_20/23nt (1.8 µM) and then subjected to crosslinking by UV. Representative figure from 3 repeats. **(E)** SPRTN-pBPA trapping assay by ssDNA. SPRTN variants (∼ 0.18 µM) carrying the E112pBPA modification were incubated in the absence or presence of FAM-labelled ssDNA_20nt (1.8 µM) and then subjected to crosslinking by UV. Representative figure from 3 repeats. **(F)** SPRTN-pBPA-substrate trapping assay by dsDNA. SPRTN variants (∼ 0.18 µM) carrying the E112pBPA modification were incubated with Cy5-N-H1 (0.5 µM) in the absence or presence of FAM-labelled dsDNA_20/23nt (1.8 µM) and subjected to crosslinking by UV. Representative figure from 3 repeats. **(G)** SPRTN-pBPA-substrate trapping assay by ssDNA. SPRTN variants (∼ 0.18 µM) carrying the E112pBPA modification were incubated with Cy5-N-H1 (0.5 µM) in the absence or presence of FAM-labelled ssDNA_20nt (1.8 µM) and subjected to crosslinking by UV. Representative figure from 3 repeats. **(H)** Schematic summary of DNA-induced conformational changes on the SPRTN N-terminal protease region. The reactions from Figure 6D-G were analysed by SDS-PAGE, followed by FITC or Cy5 scanning on an iBright 1500 imaging system (Invitrogen). SDS-PAGE was then subjected to WB using Anti-SPRTN. See also Supplementary Figure 6

Interestingly, although the crystal structure of SprT-ssDNA (PDB: 6MDX) indicates that SprT is a dimer, our results showed that only a tiny fraction of SprT dimers are present in the absence of dsDNA, as observed by native MS (Supplementary Fig. 6A). Importantly, dimeric SprT cannot be detected at all in the presence of dsDNA (Supplementary Fig. 6A), suggesting that the dimeric state seen in the crystal structure is likely a crystallographic artefact. The dimeric species of SprT-BR-E112Q was not detected in the absence of dsDNA (Fig. 6A), and only a small population of dimeric SprT-BR-E112Q bound to one molecule of dsDNA was observed (Fig. 6A). Most SprT-BR-E112Q formed a complex with dsDNA in a 1:1 ratio (Fig. 6A). SEC-MALS further confirmed that SprT-BR-E112Q did not dimerise on its own, and that only the SprT-BR-E112Q-DNA complex at a 1:1 ratio was clearly observed, with no detectable DNA-bound dimer species (Supplementary Fig. 6C). SEC-MALS analysis also excluded the possibility that ZBD-BR or ZBD form dimers on their own (Supplementary Fig. 6C). Collectively, these data suggest that dsDNA does not induce SPRTN dimerisation.

Based on the increased solvent-accessible surface area detected by native MS, we built a SprT-BR-E112Q-dsDNA model with a more open catalytic groove (Fig. 6C, Supplementary Fig. 6D), in which the key catalytic residue E112 is well exposed to solvent. To further experimentally support this model, we directly targeted E112 by bioorthogonal engineering. 2D-HSQC spectra showed similar profiles for the two constructs, SprT and SprT-BR-E112Q, indicating that they share a similar overall fold^18^ and that the mutation at E112 does not impair structural integrity. We then took advantage of this and introduced the unnatural amino acid pBPA at residue E112 in all three SPRTN constructs (SprT, SprT-BR, and BR-SprT) to assess UV-mediated trapping efficiency toward surrounding substrates (i.e., SPRTN itself or the model substrate H1), thereby probing the openness of the catalytic groove. As expected, only the SprT-BR-E112pBPA construct showed an increase in UV-induced crosslinked species (primarily dimers) upon the addition of dsDNA, indicating that the catalytic groove of one SprT-BR-E112pBPA molecule engages at least one other molecule (Fig. 6D, lane 10, Supplementary Fig. 6E). In contrast, no trapped species were detected for either SprT-E112pBPA or BR-SprT-E112pBPA, suggesting that their catalytic grooves cannot be properly opened by dsDNA (Fig. 6D, lanes 8 and 12). A heat-denatured SprT-E112pBPA sample in the absence of DNA also displayed strong UV-induced crosslinking, further confirming that native SprT adopts a compact fold (Supplementary Fig. 6F). Similarly, ssDNA induced a trapping effect on SprT-BR-E112pBPA, but not on the other two constructs (Fig. 6E). These data suggest that the structure of SprT in complex with ssDNA (PDB: 6MDX) remains in a closed conformation.

The trapping effect was more pronounced when H1 was used as the substrate (Fig. 6F, G). In the presence of DNA and under UV crosslinking, a large amount of crosslinked H1 species accumulated at the top of the SDS-PAGE gel in the SprT-BR-E112pBPA condition, whereas the other two SPRTN constructs showed either no detectable or only a minor trapping effect (Fig. 6F, G). This indicates that the native substrate, H1, can access the catalytic centre of SprT-BR in a DNA-dependent manner. Similar to H1, M1-diUb was also crosslinked to SprT-BR-E112pBPA, but not to the other two constructs (Supplementary Fig. 6G).

The DNA-induced conformational change and DNA-engaged substrate trapping suggest that SPRTN activation by DNA, and the subsequent substrate proteolysis, occur in a highly dynamic and transient manner. Our experimental data demonstrate a complex mechanism of DNA-induced conformational changes within the N-terminus of SPRTN, which contains the catalytic and DNA-binding elements. On this basis, we propose a working model (Fig. 6H). Although the isolated MPD exhibits DNA-independent protease activity, steric interference between the MPD and ZBD constrains SprT in an auto-inhibited state that cannot be released by DNA alone. ZBD, BR, and DNA act together, in trinity, to induce an open conformation of the SPRTN N-terminus, thereby relieving auto-inhibition *in cis*, exposing the catalytic groove to substrates, and enabling optimal proteolysis while facilitating the Ub-activation effect.

## Discussion

### FRET assay: a powerful tool to monitor SPRTN kinetics in real time

Our understanding of the regulation of SPRTN’s enzymatic activity has gradually grown over the past decade. Multiple biochemical characterization has greatly expanded the knowledge of both SPRTN auto-cleavage and substrate cleavage.

H1 has long been used as a model substrate to characterise SPRTN proteolysis *in vitro*^8,15,16,18^. To probe the cleavage products, sensitivity has been improved from low-resolution Western blotting^15^ to a higher-resolution fluorescent-labelling method^16^. However, apart from mapping cleavage sites in core histones^7^, major cleavage sites on H1 have not been identified. For the first time, the identification of H1 cleavage sites by intact mass analysis in this work enabled us to further develop a short peptide as a powerful and ideal substrate to monitor SPRTN kinetics in real time by FRET. With the additional help of long Ub chains, detecting SPRTN activity has never been so sensitive and straightforward (Fig. 1). Based on this method, we further validated other inhibitory factors, including 1,10-phen and Zn^2+^. It also helped us to quantitatively compare auto-inhibition and reactivation across different SPRTN truncations (Fig. 4). In addition, using 96-well plates and a plate reader equipped with basic filter sets to detect the FAM signal, this method can be easily adapted to a high-throughput format for inhibitor screening. Being simple, rapid and highly sensitive, the FRET assay has provided a powerful and promising tool for wider application in drug discovery, and therapeutics centred on SPRTN.

### SPRTN auto-cleavage: suicide or highly substrate-selective?

An “autocatalytic off-switch” model proposed that SPRTN auto-cleavage results in its inactivation^8^. A subsequent study further showed that mono-ubiquitination of SPRTN leads to direct inactivation by promoting auto-cleavage *in trans*^17^. However, it has also been reported that auto-cleaved SPRTN retains an enzymatic activity level similar to SPRTN-ΔC^8^. Thus, the notion of “inactivation” is contradictory in the context of SPRTN proteolysis.

In this study, although the residual protease activity of auto-cleaved SPRTN is very low, we found that the Ub activation and Ub avidity effects are still present. The activity of auto-cleaved SPRTN can be re-gained by longer Ub chains, making it highly selective toward the model ubiquitinated substrate (Fig. 2). This strongly suggests that auto-cleaved SPRTN predominantly targets DPCs modified with higher levels of ubiquitination. Together with its high stability, auto-cleaved SPRTN could, in fact, exert a long-term effect on substrate cleavage, rather than being inaccurately considered “inactive.”

Therefore, we propose that SPRTN is only partially inactive upon auto-cleavage. Auto-cleavage undoubtedly causes the loss of C-terminal regulatory modules, including the p97- and PCNA-interacting motifs and nuclear localisation signals. However, the high stability and remaining activity of the N-terminal fragment allow SPRTN to continue substrate proteolysis with high selectivity via the Ub avidity effect.

### SprT auto-inhibition and release: fine-tuning protease activity via conformational change

It has long been unclear how DNA activates SPRTN. The protease and DNA-binding properties of SPRTN, along with other C-terminal regulatory elements, have made it challenging to dissect its multiple, complex molecular mechanisms. Because of the high instability of the MPD protease domain, solving its structure by either solution NMR or crystallography has been difficult, and a detailed biochemical characterisation of MPD has therefore been absent. In this study, we successfully increased the stability of MPD, allowing us to further demonstrate that the isolated MPD domain retains basal intrinsic protease activity. Most importantly, MPD is neither DNA-dependent nor Ub-avidity-dependent (Fig. 3).

It was recently proposed that ubiquitin stabilises an open conformation of SPRTN based on MD simulations using a ColabFold-predicted SprT structure pre-configured in an open conformation as the starting model^21^. However, the effect of DNA was not considered in these simulations, nor was the dynamic transition from a compact to an open conformation induced by DNA captured. Our work directly addresses these gaps. Firstly, we have experimentally demonstrated that ubiquitin does not activate SPRTN in the absence of DNA, for both full-length SPRTN^16^ and SprT-BR (Supplementary Fig. 1D, E), indicating that ubiquitin alone, although capable of interacting with SPRTN, is not sufficient to exert an allosteric effect on SPRTN when DNA is absent. Thus, DNA is the primary and indispensable factor for SPRTN activation. Secondly, we identified an auto-inhibitory mechanism within the SprT construct mediated by a steric effect between the MPD and ZBD (Fig. 4). Finally, we show that DNA activates SPRTN by opening the inhibited protease region (Fig. 6), thereby enabling subsequent stabilisation of this open state by ubiquitin. The BR must be intramolecularly positioned close to the protease region. However, in the absence of DNA, BR alone cannot induce the open conformation, further underscoring the leading role of DNA in SPRTN activation. ZBD, BR, and DNA must act cooperatively to form a complex to allosterically induce the open conformation (Fig. 5). The opening of the compact protease region relieves the intramolecular auto-inhibition.

Altogether, this work resolves a major outstanding question in the SPRTN field: the detailed molecular mechanism by which DNA activates SPRTN proteolysis, a problem that has remained unclear for a long time. In summary, SPRTN employs an exquisitely complex regulatory mechanism that allows precise tuning of its activity at every stage of DPC repair, from initial DNA-induced activation to the rapid proteolysis of ubiquitinated DPC substrates.

Although our pBPA-trapping method has provided promising insights into the DNA-induced dynamics of SPRTN, it is still important to determine the structures of the SPRTN protease region in the absence and presence of DNA to directly visualise the allosteric transition from auto-inhibition to activation. The structure of the ZBD-BR-DNA complex would also help clarify the detailed mechanism by which these three key factors regulate the openness of SPRTN.

## Methods

### Protein expression and purification

#### H1 and Ub purification

His-tagged H1 variants (including M1-Ub4-H1), His-tagged monoUb, M1-linked Ub chains, and N^15^-labelled His-tagged monoUb were all expressed and purified following the previously reported method^16^.

#### SPRTN purification

SPRTN-FL was expressed and purified following the previously reported method^15^. GST-SPRTN-ΔC, tag-free SPRTN truncations (including 1-227, MIU-SprT, SprT, BR-SprT, SprT-BR, SprT-BR-E112Q, ZBD-BR and ZBD) were all expressed and purified following the previously reported method^16^.

To express the E112pBPA variants of SprT, SprT-BR and BR-SprT, *E.coli* BL21(DE3) strain carrying both the SPRTN variant plasmid and pEVOL-pBpF (Addgene: 31190) was grown in LB medium (containing 100 µM ZnCl2) at 37 °C with constant shaking until OD600 reached to ∼0.8. Each 1 L culture was then induced by adding 0.5 mL 1M IPTG, 1 mL 1M pBPA (prepared in 1M NaOH) ^23^ and 2g L-arabinose (powder), making the final concentration of IPTG at 0.5 mM, pBPA at 1mM and L-arabinose at 0.2%. Incubate at 18 °C overnight. The purification procedure follows the same as SprT-BR from the previously reported method^16^.

To express GST-MPD and L99A variant, *E. coli* BL21(DE3) strain carrying the pGEX-6P1-SPRTN-MPD-WT or L99A plasmid was grown in M9 medium (containing 6 g/L Na2HPO4, 3 g/L KH2PO4, 0.5 g/L NaCl, 2 g/L D-glucose, 0.7 g/L NH4Cl, 10 mL/L100× Gibco^®^ MEM vitamin solution, 10 µM FeSO4, 10 µM CaCl2, 2 mM MgSO4, 100 µM ZnCl2, pH 7.4) at 37°C with constant shaking until OD600 reached to ∼1.5. It was induced by IPTG at a final concentration of 0.2 mM at 20 °C overnight. The harvested cell pellet was resuspended in ∼50 mL GST High Salt Buffer (100 mM Tris-HCl, pH7.4, 500 mM NaCl, 1 mM DTT) containing 0.2 mM phenylmethanesulfonyl fluoride (PMSF) and one tablet of EDTA-free protease inhibitor cocktail (Roche) and then processed by sonication. The lysate was centrifuged at 20K rpm for 30 min. The supernatant was loaded onto the home-packed column with 10 ml of Glutathione Sepharose High Performance (Cytiva) pre-equilibrated with GST High Salt Buffer. The column was further cleaned with GST Low Salt Buffer (100 mM Tris-HCl, pH7.4, 150 mM NaCl, 1 mM DTT). The protein was eluted with GST Low Salt Buffer containing 20 mM reduced GSH. The GST tag was not removed.

### Cy5 labelling

Naked H1 (S191C or M1-Cys-T2 variants) and M1-Ub4-H1 (M1-Cys-T2) were labelled with Cy5 following the previously reported method^16^.

To label SPRTN-ZBD (151-215) with Cy5, 100 µL SPRTN-ZBD (1756.7 µM) was mixed with 10 µL Cy5 (Cytiva, one vial was dissolved in 50 µL DMSO) and 50 µL Labelling Buffer (50 mM HEPES, pH 7.4, 150 mM NaCl, 0.5 mM TCEP-HCl). Incubate overnight in the dark at 4 °C. The labelled protein was separated from the free dye by dialysis against ∼ 250 mL Labelling Buffer using a 3.5K MWCO Slide-A-Lyzer Dialysis Cassette (3 mL, Thermo) at 4 °C. After ∼ 3h, the outer buffer was replaced with fresh 300 mL Labelling Buffer and dialysis was left overnight at 4 °C. The dialyzed protein was further concentrated using a 3K Amicon^®^ Ultra Centrifugal Filter (0.5 mL, Millipore) to desired concentration as the final product.

To label M1-diUb with Cy5, 100 µL M1-diUb (G152C) (2.23 mM) was mixed with 10 µL Cy5 and 100 µL Labelling Buffer. Incubate overnight in the dark at 4 °C. It was further buffer-exchanged with Labelling Buffer several times using a 3K Amicon^®^ Ultra- 15 Centrifugal Filters (0.5 mL, Millipore) until the flowthrough doesn’t show any visible blue colour from the free Cy5.

### Making dsDNAs

All the ssDNAs were diluted by DEPC-treated water to a final concentration of 100 µM respectively. To make dsDNA, equal volume of ssDNA was mixed together and incubated at 95 °C for 10 min on a heating block. The mixture was then left on the heating block switched off to allow cooling down with gradient temperature drop naturally until equilibrium to room temperature. dsDNA_20nt was made with Turner-20bp-F and Turner-20bp-R; dsDNA_20/23nt was made with Turner-20bp-F and WS-23bp-R. FAM-dsDNA_20/23nt was made with FAM-Turner-20bp-F and WS-23bp-R.

### SPRTN single-turnover cleavage assay

For the SPRTN cleavage towards H1 substrates, recombinant SPRTN (or variants) (2 µM) were incubated with H1 (1 µM) in the absence or presence of dsDNA_20/23nt (2.7 µM) in combination with Ubs (2 µM) and inhibitors (1,10-phen or ZnCl2, concentration specified in corresponding figure legend) for the indicated time at 30°C. Samples were resolved by SDS-PAGE followed by Cy5- or FAM-scanning. Details can be found in each corresponding figure legend.

### SPRTN multi-turnover FRET assay

For the SPRTN cleavage towards the model H1c peptide, recombinant SPRTN (or variants) (2 µM) were incubated with H1c peptide (20 µM) in the absence or presence of dsDNA_20/23nt (2.7 µM) in combination with Ubs (2 µM) and inhibitors (1,10-phen or ZnCl2, concentration specified in corresponding figure legend). The assay was monitored by a platereader (FLUOSTAR, BMG) with the florescence mode at Ex. 485 nm and Em. 520 nm at 30°C for 20 min. Details can be found in each corresponding figure legend.

### SPRTN-pBPA trapping assay

SPRTN variants (∼ 0.18 µM) carrying the E112pBPA modification was incubated in the absence or presence of FAM-labelled ssDNA_20nt or dsDNA_20/23nt (1.8 µM) in combination with Cy5-N-H1 or Cy5-M1-diUb (0.5 µM) at room temperature for 10 min. It was then subjected to crosslinking by UV at room temperature for 20 min. Samples were resolved by SDS-PAGE followed by Cy5- or FAM-scanning, then subjected to western blot by Anti-SPRTN.

### Microscale Thermophoresis (MST)

To determine the affinity between BR peptide and Cy5-ZBD, a serial dilution of BR peptide (3.73-61127 µM, dissolved in MST Buffer containing 50 mM HEPES, 150 mM NaCl, 0.5 mM TCEP-HCl) was mixed with Cy5-ZBD (concentration fixed at 7.5 µM). Each sample was loaded to the capillary (Nanotemper) for the measurement with 20% MST Power.

To determine the affinity between BR peptide and FAM-labelled DNAs, a serial dilution of BR peptide (prepared the same as above) was mixed with FAM-labelled DNA (concentration fixed at 0.35 µM). For the measurement with FAM-ssDNA_20nt, BR peptide at a range of 3.73-15281.8 µM was used. For the measurement with FAM-dsDNA_20nt, BR peptide at a range of 3.73-30563.5 µM was used. For the measurement with FAM-dsDNA_20/23nt, BR peptide at a range of 3.73-61127 µM was used. Each sample was loaded to the capillary (Nanotemper) for the measurement with 20% or 40% MST Power.

All the MST measurements were performed on a microscale thermophoresis instrument (Monolith NT.115, Nanotemper). The data were analysed and plotted by the MO.Affinity Analysis Software (v2.3, Nanotemper).

### Isothermal Titration Calorimetry (ITC)

Protein and DNA samples were dialysed against ITC buffer (50 mM HEPES pH 7.4, 50 mM NaCl, 0.5 mM TCEP-HCl) overnight at 4°C using the Mini Dialysis Kit (1 KDa cut-off, Cytiva) before measurements.

All the ITC measurements were conducted in a MicroCal PEAQ-ITC calorimeter (Malvern) with a standard 13-injection titration program (1st injection: 0.4 μl/injection, duration: 0.8s, spacing time: 150 s; the rest of the injections: 3 μL/injection, duration: 6s, spacing time: 150 s) at 25 °C, with a reference power at 10 μCal/s. Stir speed: 750 rpm. Initial delay: 60s. The DNAs were kept in the syringe all the time while the proteins were kept in the cell. Detailed optimized concentrations for the titration are as follows: 300 μM dsDNA_20/23nt into 40 μM SprT-BR-E112Q; 300 μM dsDNA_20/23nt into 40 μM SprT; 150 μM dsDNA_20/23nt into 100 μM ZBD-BR; 250 μM dsDNA_20/23nt into 100 μM ZBD; 300 μM ssDNA_20nt into 40 μM SprT; 300 μM ssDNA_20nt into 50 μM ZBD-BR; 100 μM ssDNA_20nt into 150 μM ZBD. The data is processed and analyzed using the MicroCal PEAQ-ITC Analysis Software with a one-site binding model.

### NMR spectroscopy

^15^N-labelled His-monoUb was prepared at a concentration of 50 or 100 uM in NMR Buffer (22 mM phosphate, pH 7.0, 55 mM NaCl, 1 mM DTT) containing 5% D2O and 0.05% NaZ. NMR spectra were recorded on a 950 MHz spectrometer (Bruker Avance III HD console) equipped with high-sensitivity 5 mm TCl cryoprobe at 25 °C.

A series of 0-4X titration pairs were made by addition of unlabelled GST-SPRTN-MPD WT or L99A or free GST to the ^15^N-labelled His-monoUb. HSQC Spectra were recorded. Spectra were initially processed by TopSpin and then analysed and plotted by MestReNova.

### Intact mass analysis (LC-MS)

Reversed-phase chromatography was performed in-line prior to mass spectrometry using an Agilent 1290 uHPLC system (Agilent Technologies inc. – Palo Alto, CA, USA). Concentrated protein samples were diluted to 0.02 mg/ml in 0.1% formic acid and 50 µl was injected on to a 2.1 mm x 12.5 mm Zorbax 5um 300SB-C3 guard column housed in a column oven set at 40 °C. The solvent system used consisted of 0.1% formic acid in ultra-high purity water (Millipore) (solvent A) and 0.1 % formic acid in methanol (LC-MS grade, Chromasolve) (solvent B). Chromatography was performed as follows: Initial conditions were 90 % A and 10 % B and a flow rate of 1.0 ml/min. A linear gradient from 10 % B to 80 % B was applied over 35 seconds. Elution then proceeded isocratically at 95 % B for 40 seconds followed by equilibration at initial conditions for a further 15 seconds. Protein intact mass was determined using a 6530 electrospray ionisation quadrupole time-of-flight mass spectrometer (Agilent Technologies Inc. – Palo Alto, CA, USA). The instrument was configured with the standard ESI source and operated in positive ion mode. The ion source was operated with the capillary voltage at 4000 V, nebulizer pressure at 60 psig, drying gas at 350 °C and drying gas flow rate at 12 L/min. The instrument ion optic voltages were as follows: fragmentor 250 V, skimmer 60 V and octopole RF 250 V. Data analysis was performed using Agilent proprietary software Masshunter Qualitative Analysis V7.0.

### Native mass spectrometry

Samples (dsDNA_20/23nt was used for the dsDNA-protein complex at 1:1 ratio) for native mass spectrometry were held on ice and desalted by buffer exchange into 50 mM ammonium acetate solution, pH 7.2 using Micro Bio-Spin™ P-6 Gel Columns (Biorad) following the manufacturer’s instructions. Protein loading was between 50 µL and 75 µL at approximately 1 mg/mL. Desalting was achieved by passing sequentially through 3 spin columns. Desalted samples were infused directly into a 6530 TOF mass spectrometer (Agilent Technologies inc. – Palo Alto, CA, USA) equipped with a standard Apollo source at a flow rate of 6 µL per minute by means of a syringe pump. Instrument source conditions were as follows: capillary 3500 V, fragmentor 380 V, desolvation gas flow 5 L/min, desolvation gas temperature 325 °C, nebuliser gas pressure 17 psi. Instrument on optic settings were: skimmer 65 V, octupole rf 750 V. The instrument was operated in positive ion mode with the detector set to 1 GHz from m/z 500 to m/z 20,000.

Data analysis was performed manually using the Masshunter Qualative Analysis V7.0. Protein deconvolution was performed using ESIprot^24^. Charge states were determined using a charge table. Oligomeric states were identified from consecutive ion series. The solvent accessible surface area (SASA^2^) was calculated from the charge states using the equation described by Hall and Robinson^25^. For dynamic surface area plots, ion intensities were normalised by division of each value by the largest for each molecular species and plotted against the calculated solvent accessible surface area (SASA^2^) fitted by Caussian – least squares fit (Prism).

### Size-exclusion chromatography coupled multiple-angle laser light scattering (SEC-MALS)

Protein samples were prepared at ∼ 2 mg/mL. 100 µL of each sample was applied to the pe-equilibrated SEC column (Superdex 75 10/300 GL, GE Healthcare) at a flow rate of 0.5mL/min using the buffer containing 50 mM HEPES pH 7.4, 150 mM NaCl, 0.5 mM TCEP-HCl. The eluted protein was monitored via online static light-scattering (DAWN HELEOS 8+, Wyatt Technology), differential refractive index (Optilab T-rEX, Wyatt Technology) and UV (SPD-20A, Shimadzu) detectors. Data were analysed by Astra 6.1 (Wyatt Technologies) with a refractive increment value of 0.185 mL/g.

### Modelling method to build SprT-BR-dsDNA

A molecular model of DNA-induced SPRTN opening structure was constructed starting from the crystallographic SPRTN-ssDNA complex (PDB: 6MDX). The model was processed to remove alternate conformations. Additional residues corresponding to the BR domain (216-227) were added from an Alphafold3 model of SPRTN 1-236 avoiding any regions with low confidence (pIDDT<50). dsDNA was positioned on this model based on a positive surface patch identified by calculating an electrostatic surface using Pymol, as well as based on the phosphate and base positions of the Cytosine 1 from the ssDNA which binds to the ZBD in the crystal structure. To model DNA-induced opening, a hinge between MPD and ZBD was selected around residue 154-156 and the model was rotated to maximize additional favourable contacts between the DNA backbone and both the BR and MPD domains.

### Data analysis

Kinetic data from SPRTN cleavage assay were fitted with one phase exponential decay - least squares fit (Prism). Significant analysis was performed by one-way ANOVA (Prism). *p <0.05; **p <0.005; ***p <0.0005. Details can be found in each corresponding figure legend.

## Data and Code Availability

This study did not generate code or reposited datasets.

## Acknowledgements

The authors thank Dr Raimundo Freire from Hospital Universitario de Canarias for kindly providing the SPRTN antibody and Dr Ben Stieglitz from Queen Mary, University of London, for kindly providing the Ub constructs. The authors thank Dr Anna Pérez- Ràfols and Prof. Yogesh Kulathu from the University of Dundee for kindly providing the branched Ub chains. The authors also thank the support from the Molecular Biophysics Suite, Department of Biochemistry as well as the Biophysics and Biochemistry Group, Centre for Medicines Discovery, University of Oxford to the access to the biophysical instruments. This work received support from the Medical Research Council programme (Grant Ref. MR/X006409/1), Breast Cancer Now (Grant Ref. 2022.11PR1570), Ministry of Education-Strat-Up Grant, Singapore (023917-00001), Toh Kian Chui Distinguished Professorship Award/LKC Medicine, Singapore, to K.R. C.R. acknowledges support for upgrading the 950 MHz spectrometer with funding from the University of Oxford Wellcome Institutional Strategic Support Fund, the John Fell Fund, the Edward Penley Abraham Cephalosporin Fund, and the Engineering and Physical Sciences Research Council (Grant Ref: EP/R029849/1). J.A.N. acknowledges support for his research from National Cancer Institute P01 CA092584 and Cancer Research UK A24759.

## Author contributions

Conceptualization: W.S. and K.R. Investigation: W.S., J.A.N., Y.Z., R.C., C.R. NMR: W.S. and C.R. MS: R.C. Writing – Original draft: W.S. and K.R. Writing – Review & Editing: W.S., J.A.N. and K.R. with input from all authors. Funding Acquisition: K.R.

## Additional information

### Supplementary inforamtion

**Correspondence** and requests for materials should be addressed to Kristijan Ramadan.

