## Supplementary Figures and materials for "DNA-induced conformational changes in SPRTN relieve its auto-inhibitory effect on protease activity"

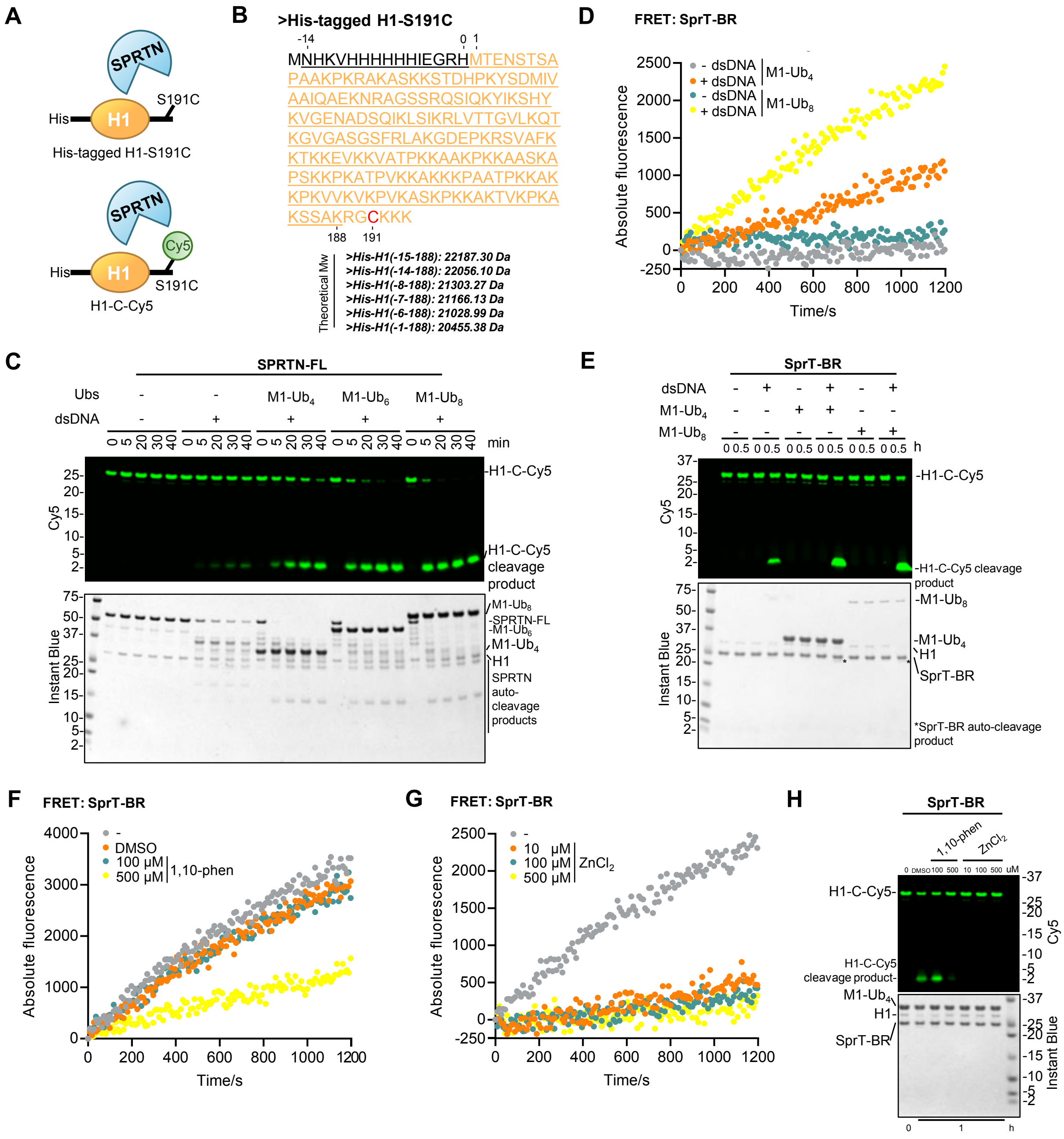


### Supplementary Figure 1

**Supplementary Figure 1. A powerful FRET assay allows for monitoring Ub-activated SPRTN kinetics in real time**

**(A)** Schematic of the His-tagged H1-S191C for the intact MS (upper) and the Cy5-labelled His-tagged H1-S191C (H1-C-Cy5) for the single-turnover SPRTN cleavage assay (bottom).

**(B)** Full protein sequence of the His-tagged H1-S191C. Residues from H1 are colored in orange. Residues from the tag are indicated in dark. The underlined sequence is the major cleavage product (His-H1 (-14-188)) according to intact MS. Theoretical Mw of other cleavage products is also indicated.

**(C)** Single-turnover SPRTN cleavage assay towards H1-C-Cy5 in the presence of M1-linked Ub chains with different lengths. Recombinant full-length SPRTN (2 μM) and H1-C-Cy5 (1 μM) were incubated with corresponding M1-linked Ub chains (2 μM) in the absence or presence of dsDNA_20/23nt (2.7 μM) for the indicated time at 30°C. Representative figure from 3 repeats.

**(D)** Multi-turnover SprT-BR cleavage assay towards H1c peptide monitored by FRET. Recombinant SprT-BR (2 μM), H1c peptide (20 μM) and M1-linked Ub chains (tetraUb or octaUb, 2 μM) were incubated in the absence or presence of dsDNA_20/23nt (2.7 μM) at 30°C.

**(E)** Single-turnover SprT-BR cleavage assay towards H1-C-Cy5. Recombinant SprT-BR (2 μM) and H1-C-Cy5 (1 μM) were incubated with different combinations of dsDNA_20/23nt (2.7 μM), M1-tetraUb (2 μM) or M1-octaUb (0.2 μM) for 30 min at 30°C. Representative figure from 3 repeats.

**(F, G)** Multi-turnover SprT-BR cleavage assay towards the H1c peptide monitored by FRET in the presence of 1,10-phenanthroline (1,10-phen) (F) or $ZnCl_2$ (G). Recombinant SprT-BR (2 μM), H1c peptide (20 μM), M1-octaUb (2 μM) and dsDNA_20/23nt (2.7 μM) were incubated in the absence or presence of DMSO, 1,10-phen or $ZnCl_2$ with the indicated concentration at 30°C.

**(H)** Single-turnover SprT-BR cleavage assay towards H1-C-Cy5 in the presence of 1,10-phen or $ZnCl_2$. Recombinant SprT-BR (2 μM), H1-C-Cy5 (1 μM), M1-octaUb (2 μM) and dsDNA_20/23nt (2.7 μM) were incubated in the absence or presence of 1,10-phen or $ZnCl_2$ with the indicated concentration for 1h at 30°C. Representative figure from 3 repeats.

The FRET signals for FAM from Figure S1D, S1F-G were monitored by a platereader (POLARSTAR, BMG) with the fluorescence mode at Ex. 485 nm and Em. 520 nm for 20 min. Each curve is averaged from 3 repeats. The reactions from Figure S1C, S1E and S1H were resolved by SDS-PAGE followed by Cy5-scanning on Typhoon FLA 9500 (GE Healthcare) and Instant Blue staining.

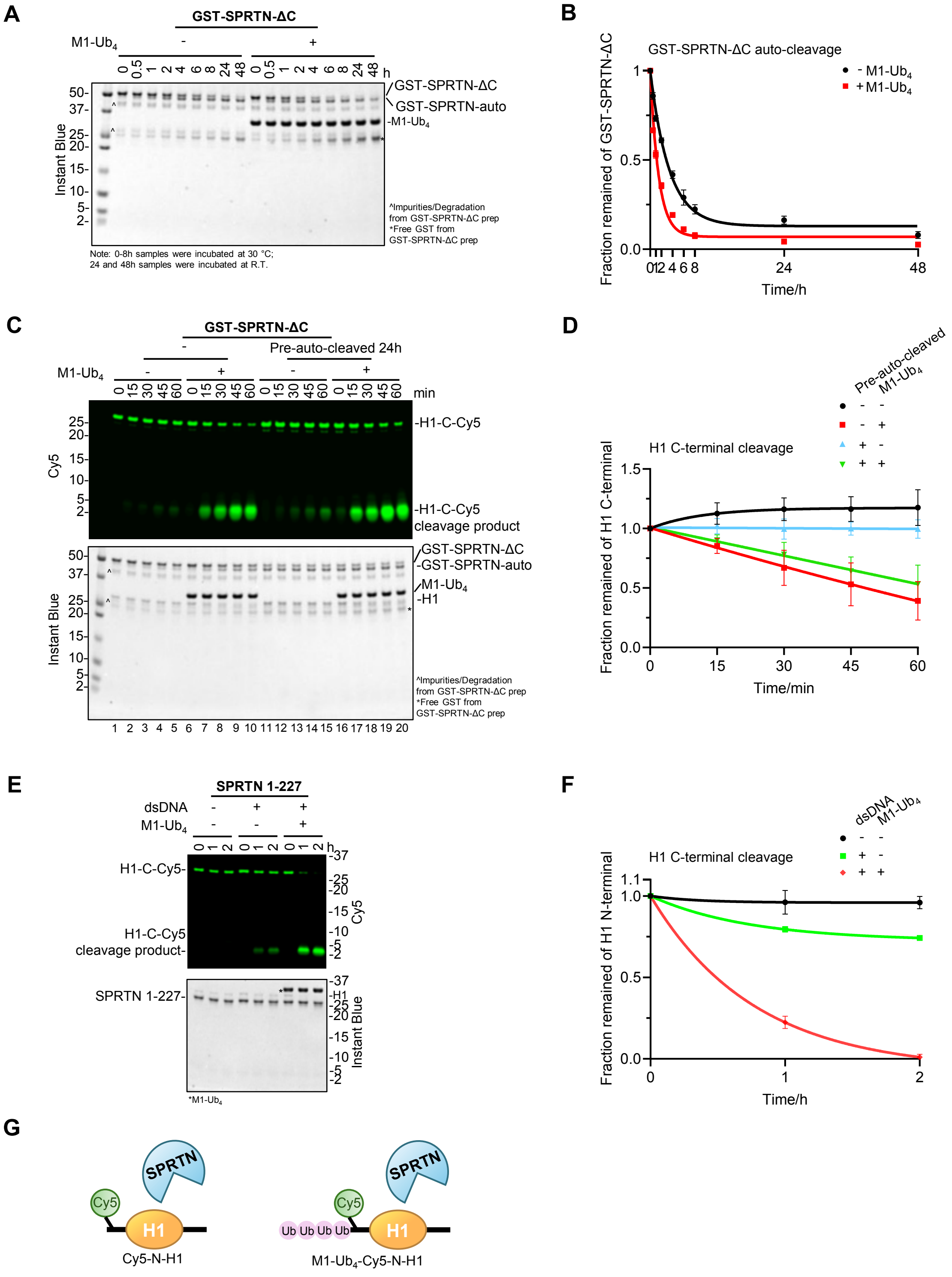


### Supplementary Figure 2

**Supplementary Figure 2. Auto-cleaved SPRTN N-terminus can be potentiated by Ub chains**

**(A)** GST-SPRTN-ΔC auto-cleavage assay. Recombinant GST-SPRTN-ΔC and dsDNA_20/23nt (2.7 μM) were incubated in the absence or presence of M1-tetraUb for the indicated time at 30°C, except that the samples for 24 and 48 h were incubated at R.T. The reaction was analysed by SDS-PAGE. Representative figure from 2 repeats.

**(B)** Auto-cleavage kinetics of GST-SPRTN-ΔC from Figure S2A. The band intensity for full-length GST-SPRTN-ΔC was quantified by the iBright Analysis Software (Invitrogen). Kinetic data were fitted with one phase exponential decay - least squares fit (Prism). n=2. Error bar, SD.

**(C)** Single-turnover GST-SPRTN-ΔC cleavage assay towards H1-C-Cy5 with or without pre-auto-cleavage. Lane 1-10: Recombinant GST-SPRTN-ΔC (2 μM), H1-C-Cy5 (1 μM) and dsDNA_20/23nt (2.7 μM) were incubated in the absence or presence of M1-tetraUb (2 μM) for the indicated time at 30°C. Lane 11-20: Recombinant GST-SPRTN-ΔC (2 μM) was pre-auto-cleaved by dsDNA_20/23nt (2.7 μM) at R.T. for 24 h and then added with H1-C-Cy5 (1 μM) for substrate cleavage in the absence or presence of M1-tetraUb (2 μM) for the indicated time at 30°C. Representative figure from 3 repeats.

**(D)** Cleavage kinetics of the signal from the full-length H1-C-Cy5 substrate (C-terminal cleavage rate) from Figure S2C.

**(E)** Single-turnover SPRTN 1-227 cleavage assay towards H1-C-Cy5. Recombinant SPRTN 1-227 (2 μM) and H1-C-Cy5 (1 μM) were incubated with different combinations of dsDNA_20/23nt (2.7 μM) and M1-tetraUb (2 μM) for the indicated time at 30°C. Representative figure from 3 repeats.

**(F)** Cleavage kinetics of the signal from the full-length H1-C-Cy5 substrate (C-terminal cleavage rate) from Figure S2E.

**(G)** Schematic of Cy5-N-H1 and M1-$Ub_4$-Cy5-N-H1 for the single-turnover SPRTN cleavage assay.

The Cy5 signals from Figure S2C and S2E were analysed by ImageJ. Kinetic data were fitted with one phase exponential decay - least squares fit (Prism). n=3. Error bar, SD.

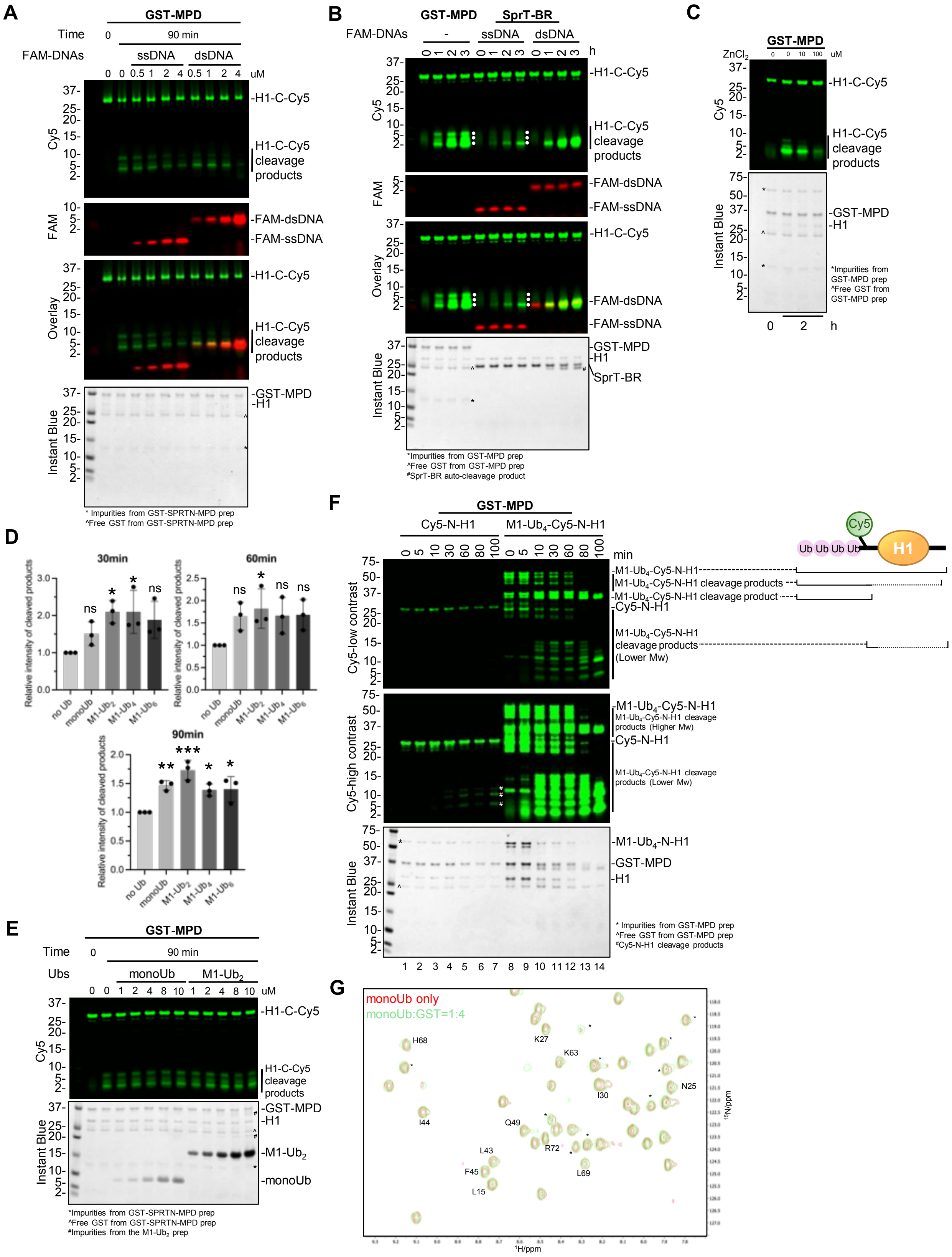


### Supplementary Figure 3

**Supplementary Figure 3. The isolated MPD is independent of both DNA activation and Ub avidity effects**

**(A)** Single-turnover GST-MPD cleavage assay towards H1-C-Cy5 in the presence of FAM-labelled DNAs. Recombinant GST-MPD (2 μM) and H1-C-Cy5 (1 μM) were incubated in the absence or presence of FAM-labelled DNAs (ssDNA_20nt or dsDNA_20/23nt) with the indicated concentration at 30°C for 90 min. Representative figure from 3 repeats.

**(B)** Activity comparison between GST-MPD and SprT-BR by single-turnover cleavage assay towards H1-C-Cy5. Recombinant GST-MPD (2 μM) and H1-C-Cy5 (1 μM) were incubated in the absence of DNA for the indicated time at 30°C. Recombinant SprT-BR (2 μM) and H1-C-Cy5 (1 μM) were incubated in the presence of FAM-labelled DNAs (ssDNA_20nt or dsDNA_20/23nt, 2.7 μM) for the indicated time at 30°C. Cleavage products are indicated with white dots. Representative figure from 3 repeats.

**(C)** Single-turnover GST-MPD cleavage assay towards H1-C-Cy5 in the presence of $ZnCl_2$. Recombinant GST-MPD (2 μM) and H1-C-Cy5 (1 μM) were incubated in the absence or presence of $ZnCl_2$ with the indicated concentration at 30°C for 2 h. DNA is not added. Representative figure from 3 repeats.

**(D)** Quantification of the H1-C-Cy5 substrate cleavage products from Figure 3B. Total Cy5 signals from H1-C-Cy5 cleavage products (region below 25 KDa) were analysed by the iBright Analysis Software (Invitrogen). Data were normalised by comparing to the condition without Ub at each time point and analysed by one-way ANOVA (Prism). N=3. Error bar, SD. *p<0.05; **p<0.005; ***p<0.0005; ns: not significant.

**(E)** Single-turnover GST-MPD cleavage assay towards H1-C-Cy5 with Ubs. Recombinant GST-MPD (2 μM) and H1-C-Cy5 (1 μM) were incubated with monoUb or M1-diUb with the indicated concentration for 90 min at 30°C. DNA is not added. Representative figure from 3 repeats.

**(F)** Single-turnover GST-MPD cleavage assay towards the model substrate Cy5-N-H1 or M1-$Ub_4$-Cy5-N-H1. Recombinant GST-MPD (2 μM) and H1 substrates (1 μM) were incubated for the indicated time at 30°C. DNA is not added. Cleavage products from Cy5-N-H1 are indicated with hash marks. Representative figure from 3 repeats.

**(G)** Superimposition of the 950 MHz $^{1}H$-$^{15}N$ HSQC spectra of monoUb alone (in red contours) and monoUb with the addition of GST at a molar ratio of 1:4 (in green contours). Spectra were plotted by MestReNova. The signals were assigned according to the reported chemical shifts of monoUb (BMRB entry: 17769). *Signals from the N-terminal His tag on monoUb.

The reactions from Figure S3A-C and S3E-F were analysed by SDS-PAGE, followed by Cy5 scanning on Typhoon FLA 9500 (GE Healthcare) and Instant Blue staining. SDS-PAGE from Figure S3A and S3B was additionally analysed by FAM scanning on Typhoon FLA 9500 (GE Healthcare).

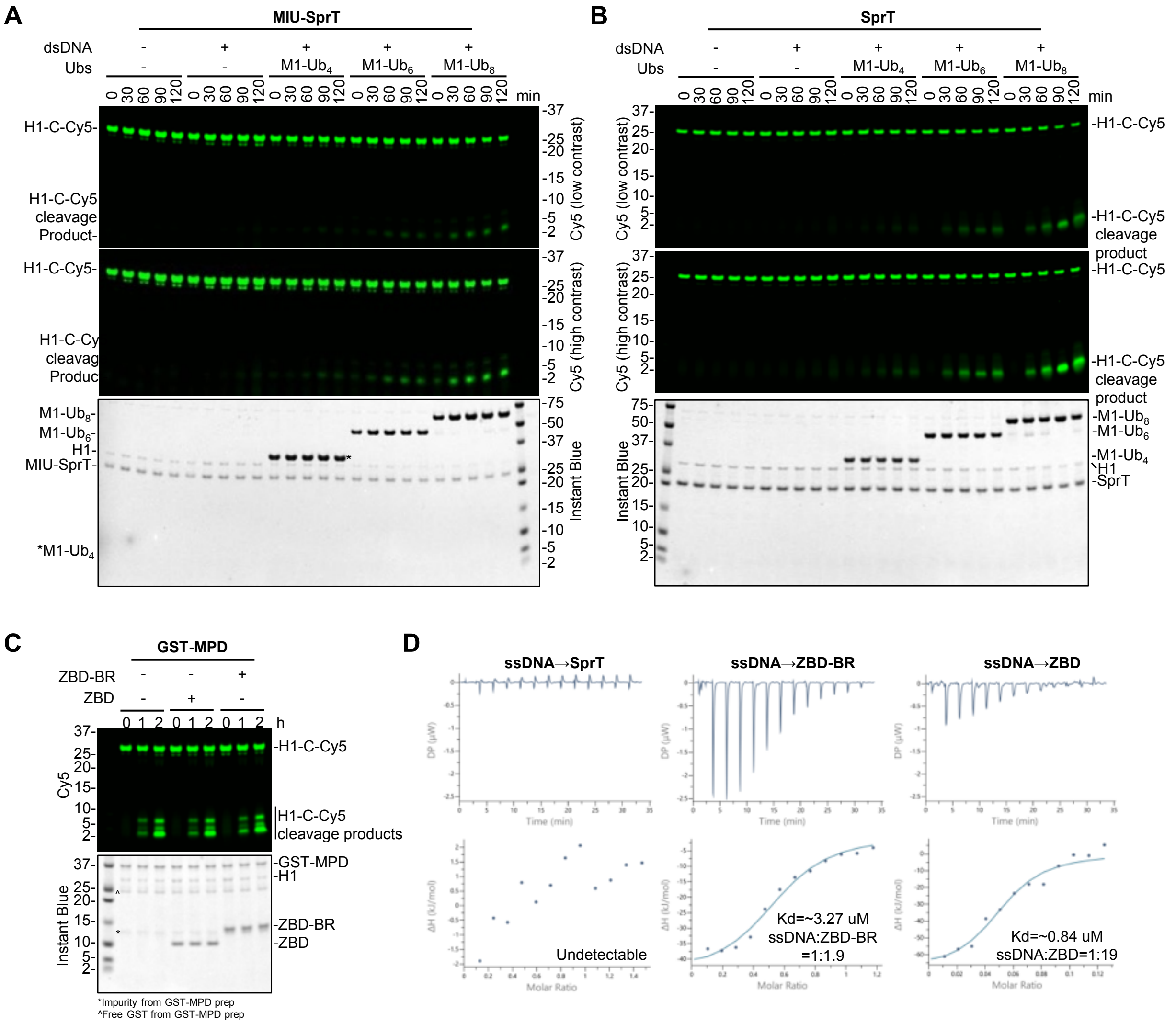


### Supplementary Figure 4

**Supplementary Figure 4. The steric effect between ZBD and MPD maintains the SPRTN N-terminal protease region in auto-inhibition**

**(A)** Single-turnover MIU-SprT cleavage assay towards H1-C-Cy5 in the presence of M1-linked Ub chains with different lengths. Recombinant MIU-SprT (2 μM) and H1-C-Cy5 (1 μM) were incubated with corresponding M1-linked Ub chains (2 μM) in the absence or presence of dsDNA_20/23nt (2.7 μM) for the indicated time at 30°C. Representative figure from 3 repeats.

**(B)** Single-turnover SprT cleavage assay towards H1-C-Cy5 in the presence of M1-linked Ub chains with different lengths. Recombinant SprT (2 μM) and H1-C-Cy5 (1 μM) were incubated with corresponding M1-linked Ub chains (2 μM) in the absence or presence of dsDNA_20/23nt (2.7 μM) for the indicated time at 30°C. Representative figure from 3 repeats.

**(C)** Single-turnover GST-MPD cleavage assay towards H1-C-Cy5 in the presence of ZBD or ZBD-BR. Recombinant GST-MPD (2 μM) and H1-C-Cy5 (1 μM) were incubated with ZBD or ZBD-BR (2 μM) for the indicated time at 30°C. DNA is not added. Representative figure from 3 repeats.

**(D)** Isothermal titration calorimetry (ITC) analysis of the interaction between ssDNA_20nt and different SPRTN N-terminal variants. The dissociation constant (Kd) and stoichiometry of binding (N) are indicated here and summarised in Table S1A.

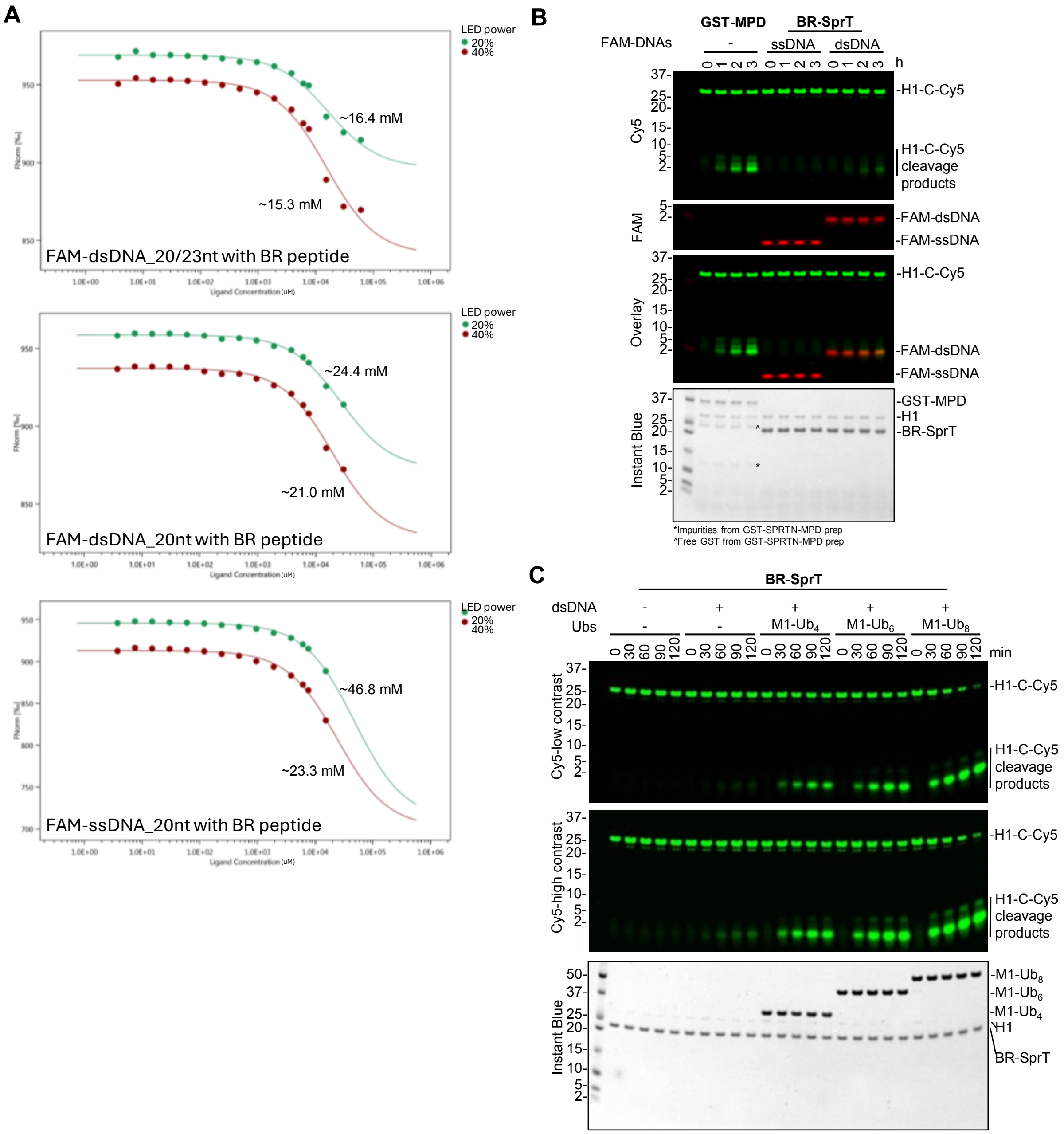


Supplementary Figure 5

**Supplementary Figure 5. The ZBD-BR-DNA trinity releases the auto-inhibition**
**(A)** MST affinity analysis between BR peptide and FAM-labelled DNAs under 20% or 40% LED power. The dissociation constant (Kd) is indicated here and summarised in Table S1B. Representative figure from 3 repeats.
**(B)** The activity comparison between GST-MPD and BR-SprT by single-turnover cleavage assay towards H1-C-Cy5. Recombinant GST-MPD (2 μM) and H1-C-Cy5 (1 μM) were incubated in the absence of DNA for the indicated time at 30°C. Recombinant BR-SprT (2 μM) and H1-C-Cy5 (1 μM) were incubated in the presence of FAM-labelled DNAs (ssDNA_20nt or dsDNA_20/23nt, 2.7 μM) for the indicated time at 30°C. Representative figure from 3 repeats.
**(C)** Single-turnover BR-SprT cleavage assay towards H1-C-Cy5 in the presence of M1-linked Ub chains with different lengths. Recombinant BR-SprT (2 μM) and H1-C-Cy5 (1 μM) were incubated with corresponding M1-linked Ub chains (2 μM) in the absence or presence of dsDNA_20/23nt (2.7 μM) for the indicated time at 30°C. Representative figure from 3 repeats.
The reactions from Figure S5B-C were analysed by SDS-PAGE, followed by Cy5 scanning on Typhoon FLA 9500 (GE Healthcare) and Instant Blue staining. SDS-PAGE from Figure S5B was additionally analysed by FAM scanning on Typhoon FLA 9500 (GE Healthcare).

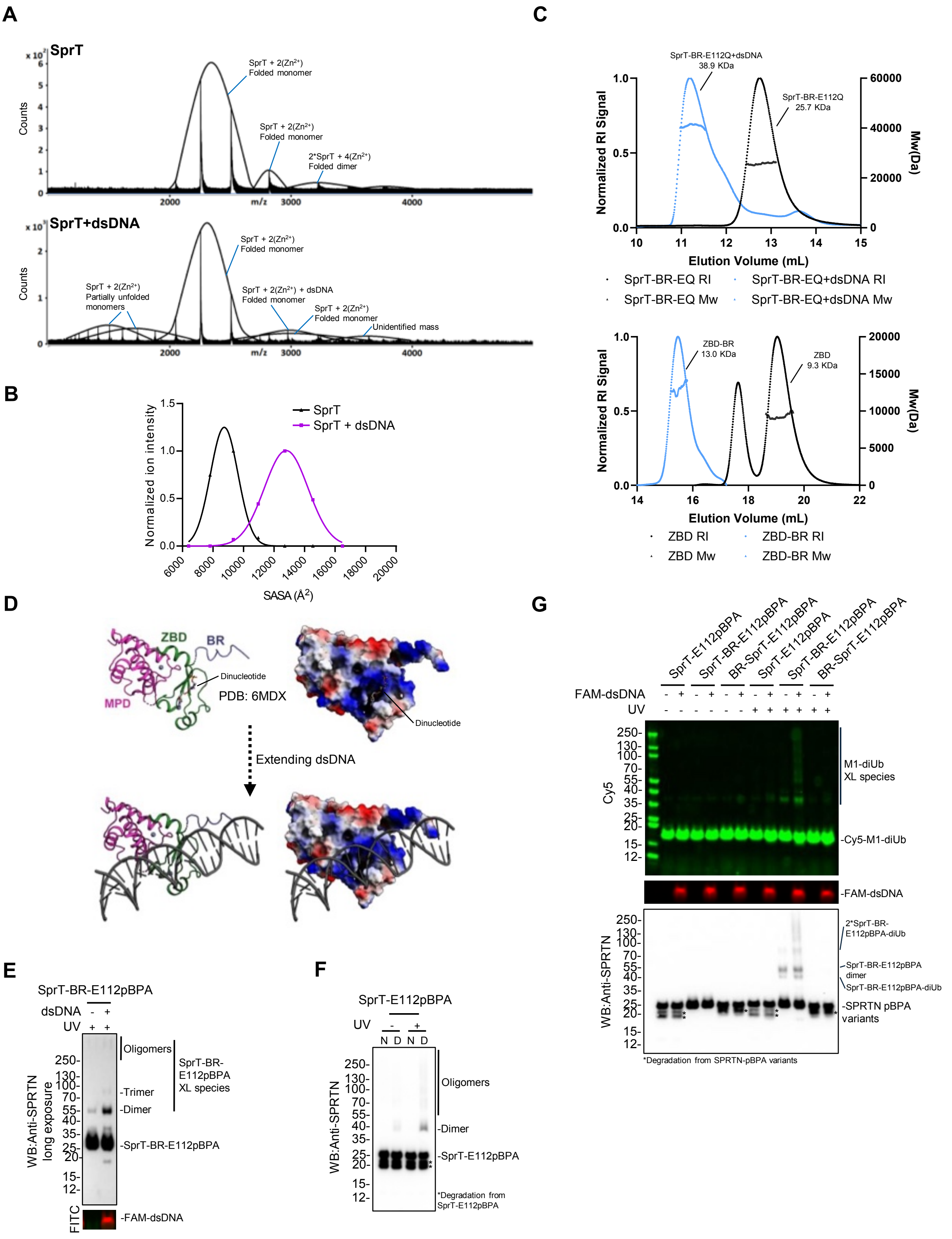


**Supplementary Figure 6**

**Supplementary Figure 6. The ZBD-BR-DNA trinity induces an open conformation of SprT-BR**

**(A)** Native MS analysis of SprT (upper) or in complex with dsDNA_20/23nt (lower).

**(B)** Gaussian distribution of the solvent accessible surface area (SASA$^2$) from Figure S6A. Data were fitted by Caussian – least squares fit (Prism).

**(C)** SEC-MALS analysis of SPRTN variants. Upper: SprT-BR-E112Q alone or in complex with dsDNA_20/23nt. Bottom: SPRTN ZBD-BR and ZBD domain. Experimental Mw is indicated respectively. Detailed parameters are listed in Table S1C. RI: refractive Index. Mw: molecular weight.

**(D)** Establishment of the starting model between SprT-BR and dsDNA based on the SprT-DNA crystal structure (PDB: 6MDX). Left: cartoon mode; Right: surface mode. The charged surface is colour-coded by red-blue spectrum (Red: negatively charged; Blue: positively charged).

**(E)** SPRTN-pBPA trapping assay by dsDNA. SprT-BR-E112pBPA (~0.18 μM) was incubated in the absence or presence of FAM-labelled dsDNA_20/23nt (1.8 μM) and then subjected to crosslinking by UV. Representative figure from 3 repeats.

**(F)** SPRTN-pBPA trapping assay with denatured SprT-E112pBPA. ~0.18 μM native or denatured SprT-E112pBPA (denatured at 60°C for 20 min) was directly subjected to crosslinking by UV. DNA is not added. The reactions were analysed by SDS-PAGE, followed by WB using Anti-SPRTN. Representative figure from 3 repeats.

**(G)** SPRTN-pBPA-Ub trapping assay by dsDNA. SPRTN variants (~0.18 μM) carrying the E112pBPA modification were incubated with Cy5-labelled M1-diUb (0.5 μM) in the absence or presence of FAM-labelled dsDNA_20/23nt (1.8 μM) and then subjected to crosslinking by UV. Representative figure from 3 repeats.

The reactions from Figure S6E and S6G were analysed by SDS-PAGE, followed by FITC or Cy5 scanning on an iBright 1500 imaging system (Invitrogen). SDS-PAGE was then subjected to WB using Anti-SPRTN.

**Table S1A: Affinity summary of SPRTN variants with DNAs from ITC (Unit: uM)**

| | | SprT-BR-E112Q | SprT | ZBD-BR | ZBD |
|---|---|---|---|---|---|
| **ssDNA_20nt** | Kd | 4.33 ± 0.98* | undetectable | 3.27 ± 0.22 | 0.84 ± 0.02 |
| | N | 0.36 ± 0.05* | N/A | 0.53 ± 0.02 | 0.05 ± 0.001 |
| **dsDNA_20/23nt** | Kd | 1.09 ± 0.19 | undetectable | 2.02 ± 0.33 | undetectable |
| | N | 0.26 ± 0.03 | N/A | 0.14 ± 0.01 | N/A |

*Data from each ITC titration pair were collected from at least 3 repeats, except for dsDNA_20/23nt with ZBD-BR and the undetectable data with 2 repeats. *Data taken from Song et al., 2025.*

**Table S1B: Affinity summary of BR peptide with ZBD or DNAs from MST**

| | LED | Kd (mM) |
|---|---|---|
| **FAM-ssDNA_20nt** | 20% | 46.75 ± 4.85 |
| | 40% | 23.27 ± 1.09 |
| **FAM-dsDNA_20nt** | 20% | 24.41 ± 2.79 |
| | 40% | 21.03 ± 1.08 |
| **FAM-dsDNA_20/23nt** | 20% | 16.38 ± 0.63 |
| | 40% | 15.33 ± 0.44 |
| **Cy5-ZBD** | 20% | 2.07 ± 0.35 |

*Data from each MST titration pair were collected from 3 repeats.*

**Table S1C: Parameters of SPRTN variants from SEC-MALS**

| | ZBD | ZBD-BR | SprT-BR-E112Q | SprT-BR-E112Q+dsDNA* |
|---|---|---|---|---|
| **Theoretical Mw (kDa)** | 8.46 | 11.70 | 25.20 | 38.30 |
| **Mn (kDa)** | 9.25 (± 3.695%) | 12.99 (± 10.685%) | 25.73 (± 1.404%) | 38.23 (± 2.002%) |
| **Mw (kDa)** | 9.26 (± 3.767%) | 13.02 (± 10.904%) | 25.74 (± 1.411%) | 38.92 (± 1.828%) |
| **Polydispersity (Mw/Mn)** | 1.001 (± 5.277%) | 1.002 (± 15.266%) | 1.000 (± 1.991%) | 1.018 (± 2.711%) |

*Mn: numeric-average molar mass; Mw: mass-average molar mass.*
**dsDNA used here is dsDNA_20/23nt (Mw:13109)*
